# Robust Phylogenetics

**DOI:** 10.1101/2025.04.01.646540

**Authors:** Qin Liu, Bui Quang Minh, Robert Lanfear, Michael A. Charleston, Shane A. Richards, Barbara R. Holland

**Author notes:** Correspondence to be sent to: School of Natural Sciences, University of Tasmania, Hobart, 7001, Australia.

## Abstract

We present a robust phylogenetic inference method, called the *trimmed log-likelihood* method, which effectively identifies fast-evolving, saturated, or erroneous sites in both simulated and empirical multiple sequence alignments. This method avoids circularity by dynamically identifying and removing sites without relying on an initial tree, allowing the specific sites removed to change as tree topology and branch lengths are estimated. Our analyses demonstrate that this method outperforms existing approaches, such as the Slow-Fast method, Tree Independent Generation of Evolutionary Rate approach, and Le Quesne Probability statistics, by removing fewer sites while still inferring phylogenies with comparable or greater accuracy. Implemented in IQ-TREE2, the trimmed log-likelihood method is user-friendly with a simple command-line interface. However, challenges remain in addressing heterogeneous evolutionary processes including compositional biases, such as GC bias. Despite these challenges, our approach offers a practical solution for improving phylogenetic inference by automatically and dynamically identifying sites to down-weight during phylogenetic analyses. We recommended that researchers compare trees inferred by varying the proportion of down-weighted sites to monitor changes in tree topology and to identify a set of candidate tree topologies for further consideration.

---

Advances in genome sequencing have created the need for improvements in phylogenetics that can keep pace with heterogeneous multi-locus data. While the incorporation of hundreds of thousands of loci holds promise for addressing complex phylogenetic challenges and mitigating stochastic errors, it is important to note that an abundance of data does not guarantee a complete solution to all aspects of evolutionary analysis (Delsuc et al., 2005; Jeffroy et al., 2006; Roger and Hug, 2006; Philippe and Roure, 2011; Philippe et al., 2011; Posada, 2016; Shen et al., 2017). Not all parts of a sequence alignment are likely to be of equal quality, and similar to any type of statistical inference, phylogenetic analysis is susceptible to the “garbage in, garbage out” principle. With the increasing volume of genomic data, systematic errors persist within the data, and may yield biased results, potentially generating strongly supported but inaccurate phylogenetic trees (Rodríguez-Ezpeleta et al., 2007; Goremykin et al., 2010; Philippe and Roure, 2011; Richards et al., 2018; Jermiin et al., 2020; Duchêne et al., 2022; Superson and Battistuzzi, 2022).

There are several types of systematic errors that can impact phylogenetic analyses, potentially resulting in incorrect or biased results. One type occurs when the assumptions of the underlying models are not met, making the model inadequate for explaining the complexity of the data (Lemmon and Moriarty, 2004; Shepherd and Klaere, 2019; Jermiin et al., 2020; Fleming et al., 2023). Another type of systematic error arises from human mistakes that affect the quality of the data, such as mislabelled sequences which may be caused by contamination, or paralogous sequences, which are genes that have diverged after duplication and may be incorrectly classified as orthologous sequences (Kozlov et al., 2016; Fleming et al., 2023).

Focusing on the first type of systematic errors, data complexity can arise from specific sites, genes, or taxa that the model struggles to account for. These sites include fast-evolving sites, which may also become saturated over time (Pisani, 2004; Shavit Grievink et al., 2013; Superson and Battistuzzi, 2022; Xia et al., 2003; Goremykin et al., 2010), as well as compositionally biased sites (Husník et al., 2011; Naser-Khdour et al., 2019; Kapli et al., 2021).

While fast-evolving sites have the potential to contain valuable phylogenetic information, they may compromise the accurate reconstruction of evolutionary relationships if multiple substitutions per site are not properly accounted for by the model (Xia et al., 2003; Cummins and McInerney, 2011; Philippe and Roure, 2011). These multiple substitutions pose a significant challenge because they can create patterns that mimic those arising from shared evolutionary history, making it difficult to distinguish true historical signals from convergent changes. This confusion can cause long branch attraction (LBA) artefacts, where unrelated sequences are erroneously grouped together as a clade (Felsenstein, 1973; Olsen, 1987; Huelsenbeck, 1997; Bergsten, 2005; Kennedy et al., 2005; Philippe et al., 2005; Goremykin et al., 2010; Fleming et al., 2023).

Another challenge related to data complexity in phylogenetic inference is the presence of compositionally biased sites. Variations in nucleotide or amino acid composition between these sites can introduce substantial complexities for phylogenetic analyses (Jermiin et al., 2004). This issue is particularly pronounced in large datasets that include thousands of loci. For instance, in vertebrate genomes, GC content can vary across chromosomal regions, with GC-biased gene conversion (gBGC), a process where G and C bases are favoured over A and T during DNA repair, leading to an accumulation of G and C bases in certain regions (Duret and Galtier, 2009; Huttener et al., 2019).

Furthermore, undetected recombination events can create the illusion of rapid evolution, complicating phylogenetic inference. Recombination, which causes different sites to follow different evolutionary histories, can make some sites appear to be fast-evolving sites, as they are fit to an incorrect tree topology (Doyle, 2022). This contributes to the “concatalescence” problem, where combining multiple genes for coalescent species tree analysis obscures the distinct evolutionary histories of individual genes (Gatesy and Springer, 2013, 2014; Doyle, 2022). Recombination adds complexity by causing different sites within a multi-gene alignment to follow different tree topologies, often mimicking fast-evolving sites (Gatesy and Springer, 2013). For example, recombination events within exons have occurred throughout mammalian history, making it challenging to interpret individual exon trees, as different parts of an exon may have distinct evolutionary histories due to these recombination events (Scornavacca and Galtier, 2017).

In addition to the inadequacies of the models in explaining the data, human mistakes also contribute to systematic errors in phylogenetic analyses. These mistakes can affect the quality of the data, such as through mislabelling sequences (Kozlov et al., 2016; Fleming et al., 2023). Mislabelling of data often occurs due to contamination during sample collection or sequence preparation, causing sequences to be incorrectly assigned to species (Merchant et al., 2014; Delmont and Eren, 2016; Breitwieser et al., 2019; Steinegger and Salzberg, 2020). Several studies have investigated the issue of contamination in empirical datasets (Merchant et al., 2014; Kozlov et al., 2016; Fleming et al., 2023). For instance, Kozlov et al. (2016) demonstrated that the proportion of mislabelled data ranges from 0.2% to 2.5% in four commonly used microbial 16S reference databases. Additionally, Merchant et al. (2014) showed in their studies that one genome, *Neisseria gonorrhoeae TCD-NG08107*, contained multiple sequences originating from the cow and sheep genomes. In addition to incorrect sequence assignments caused by contamination or mislabelling, gene misidentification, such as mistaking paralogs for orthologs, also represents a source of error. For example, Brown and Thomson (2017) showed that identifying and removing two genes containing paralogs from a dataset of 248 genes fundamentally changed the conclusions of phylogenetic analyses.

Various methods have been developed to mitigate the influence of fast-evolving or apparently fast-evolving sites on phylogenetic inference (Farris, 1969; Kuhner and Felsenstein, 1994; Brinkmann and Philippe, 1999; Hansmann and Martin, 2000; Schmidt et al., 2002; Pisani, 2004; Cummins and McInerney, 2011). In general, these methods seek to improve phylogenetic accuracy by first identifying and then removing the fastest evolving sites from a given alignment. These methods include both tree-dependent and tree-independent approaches. While many techniques are available, this study will focus on the Slow-Fast (SF), Le Quesne Probability (LQP) statistic, Tree Independent Generation of Evolutionary Rates (TIGER), and Observed Variability (OV) methods, which have been widely used to address challenges of identifying fast-evolving sites or saturated sites.

The SF method, introduced by Brinkmann and Philippe (1999), is a tree-dependant parsimony approach that identifies fast-evolving sites by quantifying the number of evolutionary steps within a predefined group using a Maximum Parsimony (MP) tree. In contrast, character compatibility scores for sites, first introduced by Quesne (1969), provide a tree-independent method for identifying fast-evolving sites. These compatibility scores assess the compatibility of a site in relation to all other sites in a multiple sequence alignment. Based on this site compatibility approach, two similar randomisation tests were developed independently by Wilkinson (1992) and Meacham (1994). These tests use the LQP statistic to evaluate whether the compatibility of a site with the other sites is no better than expected by chance, thereby identifying fast-evolving sites by high incompatibility scores. Additionally, based on similar approach to the LQP test, Cummins and McInerney (2011) introduced the TIGER method, which analyses similarity within sites. TIGER also uses site compatibility as a proxy for evolutionary rate, identifying sites with reduced compatibility as fast-evolving sites. Another similar method to TIGER is OV (Goremykin et al., 2010), which calculates the variability of each site independently by comparing the number of pairwise character-state matches to mismatches among an alignment for which the site is scored. TIGER and OV are also known as subsampling methods since they divide alignments into smaller groups of sites based on estimated evolutionary rates (Simmons and Gatesy, 2016).

Various studies have applied these techniques to identify fast-evolving sites, demonstrating both their strengths and limitations (Simmons and Gatesy, 2016; Song et al., 2016; Qu et al., 2017; Aouad et al., 2018; Klimov et al., 2018; Chang et al., 2021; Mongiardino Koch and Thompson, 2021; Aouad et al., 2022; Duchêne et al., 2022; Superson and Battistuzzi, 2022). The use of SF, LQP, TIGER, and OV approaches has proven beneficial in constructing or investigating deep phylogenetic relationships in various groups such as *Archaea*, *Pancrustacean*, *Halobacteria*, *Angiosperms*, *Sponge* and *Cupressaceae* (Rota-Stabelli et al., 2013; Qu et al., 2017; Sperling et al., 2009; Aouad et al., 2022; Superson and Battistuzzi, 2022). By using these methods to exclude fast-evolving sites, researchers have been able to explore support for different hypotheses through site subsampling (Liu et al., 2014; Superson and Battistuzzi, 2022), clarify deep phylogenetic relationships (Chang et al., 2021; Aouad et al., 2022), mitigate the LBA effect (Rota-Stabelli et al., 2013) and reduce misleading information caused by fast-evolving sites (Sperling et al., 2009).

However, some studies have shown that removing fast-evolving sites using these methods can have the opposite effect, leading to a significant reduction in phylogenetic signal (Sharma et al., 2015), resulting in poorly resolved phylogenetic trees (Cavalier-Smith et al., 2014; Klimov et al., 2018; Uribe et al., 2019; Aygoren Uluer et al., 2020). Additionally, Simmons and Gatesy (2016) argued that methods like OV and TIGER tend to favour sites with highly asymmetrical distributions of character states over those with more symmetrical distributions. This preference can cause branch length shrinkage in the internal parts of the tree, potentially increasing the risk of LBA artefacts and resulting in incorrect phylogenetic inference (Simmons and Gatesy, 2016). Furthermore, some methods, particularly the tree-dependent ones, may be circular because the identification of fast-evolving sites depends on the tree. If the initial tree used for evaluation is inaccurate, it can create a circular loop where the estimated quality of the sites influences the tree inference and vice versa.

In statistical inference a method is said to be robust if it is not overly effected by outliers or minor model violations (Tukey, 1960; Hampel, 1968; Huber, 1992). For example, in a regression setting, trimmed least squares is a robust version of regression that minimises the sum of squares for some fraction of the smallest residuals rather than all the residuals (De Menezes et al., 2021). In the context of maximum likelihood phylogenetics, this idea can be applied by considering the sum of site log-likelihood values for some fraction of sites rather than for all sites. In this sense, methods that seek to remove fast-evolving sites can all be considered as attempts at developing robust phylogenetic methods.

In this study, we introduce a robust method for identifying fast-evolving sites and other erroneous sites, known as the *trimmed log-likelihood* method, which is analogous to trimmed least squares. This approach is designed to provide reliable results in the presence of outlier sites. Outliers can signal potential errors in the data, such as the mislabelling of gene sequences for species or the incorrect inclusion of paralogs in a dataset (Brown and Thomson, 2017). Identifying outliers is valuable not only for improving data accuracy but also for uncovering significant biological processes. For example, outliers in the data may be due to positive selection, where some genomic regions have acquired convergent evolutionary changes due to environmental pressures (Blomberg and Garland Jr, 2002; Kosiol et al., 2006; Moran et al., 2008; Chang and Duda Jr, 2012; Baker et al., 2016). Additionally, outliers may result from some genes following different evolutionary histories to others, such as the case of two genes containing previously unrecognised paralogs identified by Brown and Thomson (2017), whose removal from a much larger dataset fundamentally altered the phylogenetic conclusions.

The trimmed log-likelihood method, built on the maximum likelihood framework, dynamically excludes low-likelihood sites during tree-space exploration, adjusting the sites removed as different tree topologies, branch lengths and model parameters are evaluated. This adaptability ensures that site exclusion is not predetermined, avoiding circularity.

Selecting sites for analysis is just as important as taxon sampling and model selection in phylogenetic studies (Shavit Grievink et al., 2013; Superson and Battistuzzi, 2022; Duchêne et al., 2022). Few studies have explicitly investigated the optimal cut-off point for site removal to ensure robust inference, with one of the earliest mentions being by Pisani (2004), who suggested that this threshold should be determined individually for each dataset. There is considerable variability in the proportion of sites that different methods recommend to remove, as highlighted by Simmons and Gatesy (2016), with thresholds ranging from as little as 0.6% (Owen et al., 2015) to as much as 76% (Feuda and Smith, 2015).

While determining the optimal cut-off point is valuable, we argue that deciding which sites to remove is even more critical. For instance, Shen et al. (2017) demonstrated that phylogenetic trees estimated from a dataset often depend on a small number of sites with a strong influence on the likelihood, raising questions about the robustness of conclusions, even from large alignments. The trimmed log-likelihood method we propose suggests which sites should be removed, but does not give exact guidance on how many.

In this study, we explore the impact of using the trimmed log-likelihood approach on both empirical and simulated datasets, aiming to address two main questions: (1) Under what conditions does the trimmed log-likelihood approach improve accuracy of the phylogenetic analysis? (2) How sensitive is phylogenetic inference to the proportion of sites excluded?

## Materials and Methods

The key idea of the trimmed log-likelihood method is to exclude a certain percentage of sites, rather than summing log-likelihoods over all sites. Users first specify a substitution model (e.g. GTR, or a specific instance of a model with fully specified rate parameters), which guides the search for the best-fit tree within the tree space. Users also indicate a percentage of sites to be excluded from the likelihood calculation. In standard maximum likelihood analysis, the parameters of the tree and the model of sequence evolution are then optimised with respect to the likelihood of observing every site in the alignment. In contrast, the trimmed log-likelihood method excludes a subset of the least likely sites during each step of the optimisation.

During the optimisation process, the sites to be excluded change dynamically with each update to the branch lengths, tree topology, and evolutionary model. Specifically, sites with the lowest log-likelihood for a given set of branch lengths and topology are identified as ‘outliers’ and excluded from the current likelihood calculation. This means that different trees may ignore different sets of sites, as the exclusion criteria are not predetermined but adapt throughout the search. This approach avoids circularity by ensuring that the exclusion of sites is based on the specific model and branch lengths currently under consideration.

The trimmed log-likelihood method is implemented in IQ-TREE2 (Minh et al., 2020) (http://www.iqtree.org). IQ-TREE2 supports robust phylogenetic inference on both nucleotide and amino-acid alignments under all models available.

Users can execute the trimmed log-likelihood method by specifying a proportion of sites to retain in the command line following the --robust-phy option. If the number of sites to be included based on the specified percentage is not a whole number, it is rounded up to the nearest whole number. For example, if a JC model is applied to an alignment with 886 sites, and 98% of the sites (886 *×* 0.98 = 868.28) are retained while 2% are disregarded, the total number of sites used in the analysis will be 869 sites. The corresponding command in IQ-TREE2 would be:

> iqtree2 -s data.phy –robust-phy 0.98 -m JC -pre 98

We assessed the effectiveness of the trimmed log-likelihood method implemented in IQ-TREE2 through three simulation studies and its application to two empirical datasets. Each simulation explored scenarios where accurate phylogenetic inference is challenging, allowing us to evaluate the impact of the trimmed log-likelihood method and understand how down-weighting different proportions of sites affects results.

The first simulation addressed data mislabelling, representing a form of erroneous sites in the sequences, such as mislabelling genes in multi-gene alignments. This simulation aimed to determine whether the trimmed log-likelihood method can effectively detect and remove mislabelled sites. The second simulation evaluated the performance of the trimmed log-likelihood method in cases where different evolutionary processes occur across various lineages. We aimed to determine whether the trimmed log-likelihood method can improve the accuracy of phylogeny reconstruction when the alignments exhibit heterogeneous evolutionary processes. Lastly, the third simulation investigated the performance of the trimmed log-likelihood method using a deep branching tree, similar to one of the studies by Cummins and McInerney (2011). This simulation reflects saturated deep phylogenetic relationships and we aimed to assess the effectiveness of the trimmed log-likelihood method in removing fast-evolving sites, and compared the results with those obtained using the TIGER method in Cummins and McInerney (2011). These simulations were called the *mislabelling simulations*, the *convergent process simulations* and the *saturation simulations*, respectively.

We also analysed two empirical datasets from the studies by Cummins and McInerney (2011) and Pisani (2004) using the trimmed log-likelihood method. The first dataset, the *Thermus* dataset, was previously analysed by Cummins and McInerney (2011) using the TIGER and the TREE-PUZZLE methods to remove fast-evolving sites. The second dataset consisted of an arthropod alignment, where Pisani (2004) employed the SF method (Brinkmann and Philippe, 1999) and LQP statistics (Wilkinson, 1992) for fast-evolving sites removal. We applied the trimmed log-likelihood method to both empirical datasets and compared the results with those from the original studies.

### Simulation Procedure

#### Mislabelling simulations

In the mislabelling simulations, our biological objective was to create scenarios mirroring multi-gene alignments, specifically those where two taxa in one gene were mislabelled. By introducing these mislabelled sites into our simulations, we aimed to evaluate the effectiveness of the trimmed log-likelihood approach in identifying and removing erroneous sites while accurately reconstructing evolutionary relationships. In addition to determining which sites to remove, this analysis explored the impact of mislabelling on the reliability of phylogenetic trees. We were also interested in understanding how the proportion of sites removed influences the inferred tree topologies.

In our simulations, we generated 400 10-taxon trees using the *TreeSim* package (version 2.4) in *R* (version 4.2.2) (Stadler, 2011; R Core Team, 2019), with a speciation rate of 0.9 and an extinction rate of 0.15. To investigate the impact of branch lengths on tree inference when varying proportions of sites were removed, we divided the branch lengths by factors of 10, 5, and 2.5. We organised these trees into three groups: Short (divided by 10), Medium (divided by 5), and Long (divided by 2.5), each containing 400 trees with distinct topologies and branch lengths.

For each tree in every group, a nucleotide alignment with 1000 sites was generated using a randomly generated GTR model in *seqgen* (Rambaut and Grass, 1997). The GTR rate matrices were drawn from a uniform distribution (0.5, 5), with base frequencies set to a minimum of 0.1. The remainder of the frequencies was drawn from a uniform distribution (0, 1) and then scaled by a factor of 0.6. This process resulted in 400 alignments per group, which were labelled as Short, Medium, and Long branch length simulations.

The average branch lengths of the 400 generating trees for the Short, Medium, and Long branch length simulations were 0.06, 0.14, and 0.28 substitutions per site, respectively. For each alignment, we swapped the last 200 sites (sites 801–1000) between two randomly selected taxa. Then, using the trimmed log-likelihood method in IQ-TREE2, we fit a GTR model to each alignment and inferred the best-fit trees. These alignments had varying proportions (0% to 20%, increasing by 1%) of sites with the lowest log-likelihood values removed. The decision to remove up to 20% of the sites was based on the objective of testing whether the trimmed log-likelihood could accurately identify erroneous sites. We expected that the erroneous sites would make up no more than 20% of the total, as many sites in the 200 base pairs swapped region are likely constant and thus have high likelihood values. Therefore, removing 20% of the lowest-likelihood sites would target more than just the erroneous sites. Finally, we compared the inferred trees to the generating trees to assess the effect of the trimmed log-likelihood method.

#### Convergent process simulations

In the convergent process simulations, we aimed to test our trimmed log-likelihood method using a branch specific model where some sequences contained higher proportions of GC-rich content sites. We generated concatenated sequences comprising 1000 sites in total. One portion of these sequences included two taxa with high GC content, simulated under two different models on a rooted 9-taxon tree (Fig. 1). The other portion of the sequences was generated under a JC model, using the same topology and branch lengths as shown in Fig. 1.

**Figure 1:**
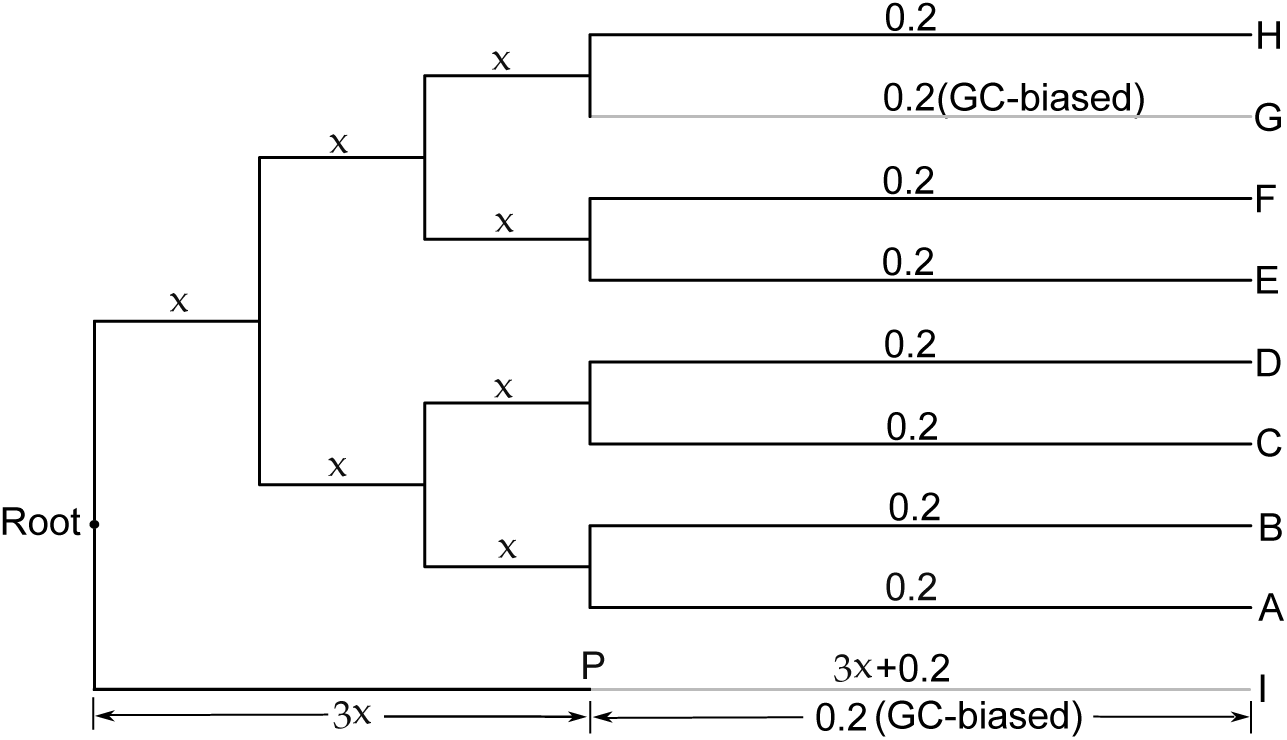
Generating tree for the convergent process simulations (branch lengths are not to scale). The internal branch length *x* varies across three simulations: 0.01, 0.03 and 0.05. GC-rich sequences were generated along the branches marked with labelled “GC-biased”, specifically, the external branch leading to taxon G and part of the branches leading to taxon I, using the UNREST model. The remaining branches used the JC model to generate sequences. At tips of taxa G and I, the GC content is around 0.8, and the AT content is around 0.2.

We conducted simulations under the same tree topology but with three different internal branch lengths: 0.01, 0.03 and 0.05 (Fig. 1). For each tree, we generated sequence alignments with varying proportions of GC-biased sites: 200 GC-biased sites with 800 JC-model sites, 300 GC-biased sites with 700 JC-model sites, and an equal split of 500 sites for each type. We labelled these simulations according to the internal branch length and the number of GC-biased sites. For example, a simulation with an internal branch length of 0.01 and 200 GC-biased sites is referred to as the “0.01-800JC-200GC” simulation. Each simulation consisted of 400 concatenated sequences. All sequence generation was performed using a custom R script (R Core Team, 2019), which is available at this GitHub repository.

For the GC-biased portion of the concatenated sequences, we used a rooted 9-taxon tree to simulate nucleotide sequences, with two branches rich in GC content, while the other branches followed a JC model (Fig. 1). Specifically, the branch leading to taxon G and part of the branch leading to the outgroup I (from point P to I) followed an unrestricted substitution model (UNREST) that generated sequences rich in GC (Yang, 1994). The UNREST model assumes a non-reversible nucleotide substitution model. The rate parameter *R* we used for the UNREST model was *R* = (*A → C, A → G, A → T, C → A, C → G, C → T, G → A, G → C, G → T, T → A, T → C, T → G*) = (5, 5, 1, 1, 5, 1, 1, 5, 1, 1, 5, 5). The average base frequencies for the GC-rich sites in taxa G and I were approximately (0.1, 0.4, 0.4, 0.1).

We then fit a JC model to all the datasets and applied the trimmed log-likelihood method to remove sites from 0% to 50%, increasing in 1% increments. The choice to remove up to 50% of sites is based on the structure of the 500JC-500GC simulations, which included 50% of sites with two taxa having a high GC content. To maintain consistency across all the simulations, we applied the same range of site removal to the other convergent simulation alignments. Our goal was to determine whether removing certain sites from the alignment improved the accuracy of the inferred tree topologies compared to the topologies inferred from the complete alignments.

#### Saturation simulations

As a negative control, we wanted to set up a simulation to assess the performance of our method under more challenging conditions. Specifically, we aimed to test how our method performs when faced with alignments containing sites that are all fairly saturated, but with no distinct class of sites that evolve by a different process - a situation where our approach is expected to struggle. The trimmed log-likelihood method is designed to identify sites with the lowest log-likelihood values, but when many of the sites in an alignment are saturated, each of them tends to have very low log-likelihoods. In such cases, it becomes difficult to determine which sites are not supporting the tree topology, as all the saturated sites have similarly low log-likelihood values, making them indistinguishable in terms of their contribution to the tree.

To simulate this, we replicated the simulations from Cummins and McInerney (2011) by generating 100 DNA alignments, each containing 999 sites using the JC model under a deep branching tree (Fig. 2a and Fig. 3a in Cummins and McInerney (2011)). This phylogenetic tree includes long terminal branches and comparatively shorter internal branches, and represents a scenario of a phylogenetic relationship that is hard to resolve.

**Figure 2:**
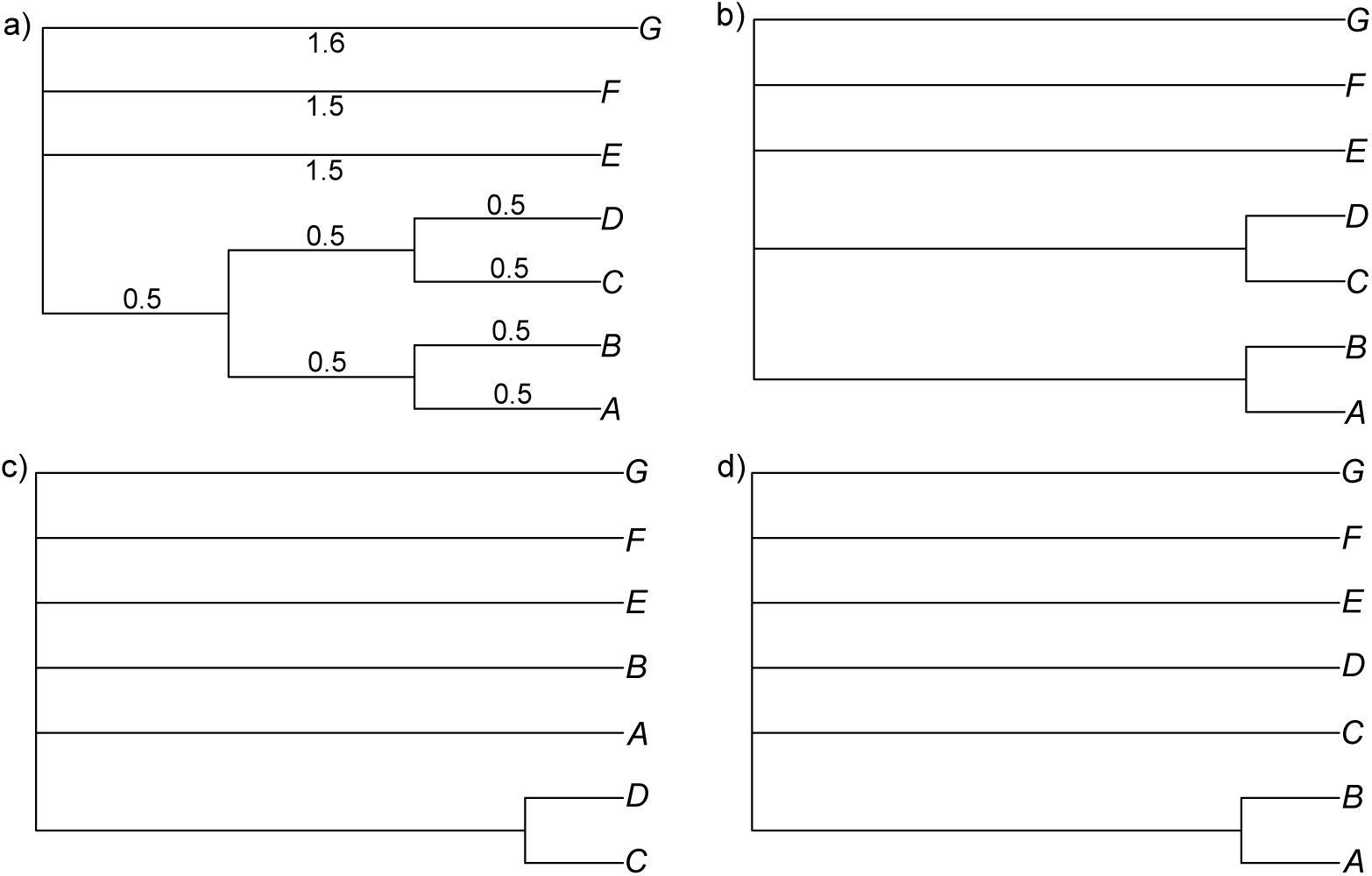
(a) The generating tree used in the saturation simulations, similar to Figure. 3a in Cummins and McInerney (2011). (b-d) Three majority-rule consensus trees constructed from 100 replicate alignments, where varying proportions of sites were removed across the simulations.

**Figure 3:**
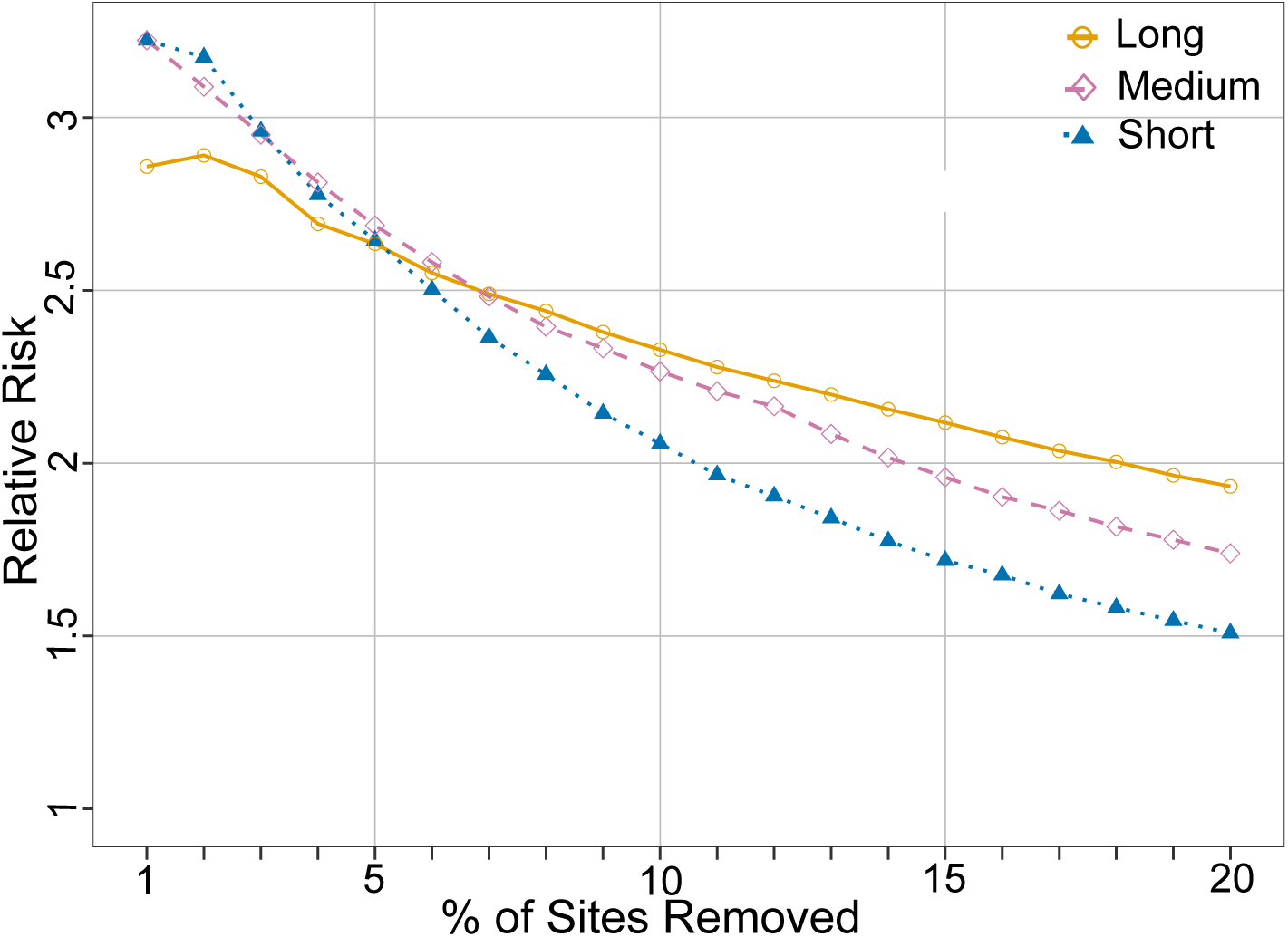
Relative risks for different proportions of removed sites for the Short, Medium and Long branch length simulations. The relative risk is the ratio of the probability of removing sites given that they were mislabelled to the probability of removing sites given that they were not mislabelled.

We then assessed the performance of the trimmed log-likelihood method by applying our method, fitting a JC model to the data and systematically removing between 0% to 50% of the sites in 1% increments. This range of site removal was based on the work of Cummins and McInerney (2011), who found that removing between 18% and 50% of sites using the TIGER method led to the best results for inferring optimal trees. For each simulation, we calculated the average RF distances between the inferred trees and the generating tree.

In Cummins and McInerney (2011), majority-rule consensus trees were constructed from trees inferred across all 100 replicate alignments, where the same proportion of sites was removed in each replicate before constructing the consensus trees. Four possible majority-rule consensus trees were identified: ((A,B),(C,D),E,F,G) (Fig. 2b), ((C,D),A,B,E,F,G) (Fig. 2c), ((A,B),C,D,E,F,G) (Fig. 2d), and the star tree. By comparing these consensus trees to the generating tree, Cummins and McInerney (2011) determined the point at which, after removing a certain number of sites, the TIGER method resulted in a consensus tree with the ((A,B),(C,D),E,F,G) topology (Fig. 2b), accurately recovering all splits in the generating tree topology except the “ABCD|EFG” split.

In our simulations, we used a similar approach. We constructed consensus trees from trees inferred from all alignments with different proportions of sites removed and compared these consensus trees to the generating tree. We also recorded the RF distances between all the inferred trees and the generating tree for each proportion of sites removed and calculated the mean RF distances across the different proportions.

### Empirical Dataset Analysis

#### Thermus dataset

We applied the trimmed log-likelihood method to the *Thermus* dataset utilised in Cummins and McInerney (2011) study. This dataset consists of an alignment of DNA sequences from the 16S rRNA gene, containing five taxa and 1273 sites. We applied three models to the dataset: GTR+G+I, TIM2+F+G4 and JC. The reasons for choosing these models are as follows: firstly, the GTR+G+I model was used in the original analysis of the *Thermus* dataset by Cummins and McInerney (2011), allowing us to directly compare our results using the trimmed log-likelihood method with those from the TIGER approach in the same study. The TIM2+F+G4 model was included because it was identified as the best-fit model by ModelFinder (Kalyaanamoorthy et al., 2017) in IQ-TREE2. Finally, the JC model, being the simplest model, was selected to evaluate whether model complexity has any impact on the results.

We systematically removed sites from 0% to 20%, with 1% increments, as we applied these three models to the data. The decision to remove up to 20% of the sites was based on the findings of Cummins and McInerney (2011), where TIGER and TREE-PUZZLE grouped the data into bins based on the evolutionary rates of the sites. The most fast-evolving bins contained 108 and 186 sites, respectively, corresponding to approximately 8.5% and 14.6% of the total sites. Therefore, we decided to remove no more than 20% of the sites in our analysis.

In the dataset, there are three thermophiles and two mesophiles. The sequences of the three thermophiles, namely *Aquifex arolicus*, *Thermotoga martima* and *Thermus aquaticus*, are richer in G and C nucleotides (around 28% for C and 36% for G), while those of the two mesophiles, *Bacillus subtilis* and *Deinococcus radiodurans*, have a relatively even distribution of nucleotide composition (around 23% for C and 32% for G) (Cummins and McInerney, 2011). Strong evidence supporting *Thermus aquaticus* and *Deinococcus radiodurans* as a sister group has been presented in previous studies (Embley et al., 1993). Following the terminology in Cummins and McInerney (2011), we refer to the tree that groups *Deinococcus radiodurans* and *Thermus aquaticus* as a clade the Reference tree, while the tree that groups *Deinococcus radiodurans* and *Bacillus subtilis* together is called the Attract tree. The term Attract tree is based on the expectation that *Deinococcus radiodurans* and *Bacillus subtilis* will be attracted to each other due to their similar GC content, a factor not accounted for in the models of sequence evolution we applied. We compared the trees inferred from alignments with various proportions of sites removed to the Reference tree and also compared our results to those in Cummins and McInerney (2011).

#### Arthropod dataset

We applied our trimmed log-likelihood method to a set of arthropod DNA sequences from the studies by Pisani (2004). These arthropod sequences are part of the Elongation Factor 1*α* gene, including 47 taxa and 866 sites. In the past, a prominent debate in the field of arthropod phylogenetics revolved around the *Mandibulata* hypothesis versus the *Myriochelata* hypothesis (Pisani et al., 2004; Pisani, 2004; Meusemann et al., 2010; Giribet and Edgecombe, 2012, 2019; Thomas et al., 2020). The *Mandibulata* hypothesis states that *Chelicerata* is monophyletic, and that *Mandibulata*, which includes the two main clades *Pancrustacea* and *Myriapoda*, is also monophyletic. Conversely, the *Myriochelata* hypothesis suggests that Myriapods should be grouped with *Chelicerates*. However, the *Mandibulata* hypothesis has consistently received more support from various studies and data, and is now widely accepted over the *Myriochelata* hypothesis (Giribet et al., 2001; Edgecombe et al., 2003; Edgecombe, 2010; Rota-Stabelli et al., 2011; Giribet and Edgecombe, 2019).

Using ModelFinder (Kalyaanamoorthy et al., 2017) implemented in IQ-TREE2, we identified the best-fit substitution model according to the AIC criterion, which was the TIM+F+I+R4 model. We fit this model to the alignments and subsequently removed from 0% to 50% of the sites in 1% increments. The rationale for testing site removal up to 50% was based on the findings of Pisani (2004), where the best results were achieved after removing 172 sites (19.9%) and 192 sites (22.6%) using the LQP and SF methods, respectively. While their analysis showed that further site removal did not yield better results, we extended the range of site removal up to 50% to explore whether removing a larger proportion of sites could improve results beyond what was observed in their study.

In their study, Pisani (2004) used the GTR+G+I substitution model to the sequences. To allow for a direct comparison, we also used this model for the data. We applied the trimmed log-likelihood method to remove from 0% to 50% of the sites in 1% increment, fitting the GTR+G+I model during each removal step. Additionally, we fit a simpler GTR model to the data and applied the trimmed log-likelihood method to remove fast-evolving sites, to evaluate the use of simpler models, as above.

The inferred trees from each removal step were compared to the strict consensus tree created from the maximum likelihood and Bayesian trees (Fig. S4 and Fig. 1 in Pisani (2004)). This consensus tree follows the *Mandibulata* hypothesis and was considered the optimal inferred tree from this arthropod dataset (Pisani, 2004).

### Performance metrics

We explored which, and how many, sites should be removed from the dataset to recover the generating tree in simulations or the reference tree in empirical studies. We also aimed to determine the optimal proportions for site removal to achieve the best phylogenetic inference. In the case of the first and second simulations, we identified whether the removed sites were mislabelled or poorly modelled. For the empirical datasets, where IQ-TREE2 gave an independent estimate of which sites were fast-evolving, we checked if these fast-evolving sites were the ones excluded.

For the simulation studies, we used Robinson-Foulds (RF) distance (Robinson and Foulds, 1981) and the Branch Score (Kuhner and Felsenstein, 1994) to measure differences in both topology and branch lengths between the inferred trees and the generating trees. These metrics allowed us to assess how well our trimmed log-likelihood method performed under different conditions by comparing the inferred trees to the known generating trees.

We also calculated the relative risk of removing erroneous or heterogeneous sites versus non-erroneous or homogeneous sites to evaluate whether our trimmed log-likelihood method effectively identified and removed problematic sites, based on the simulation settings. For example, the relative risk of removing mislabelled versus non-mislabelled sites, at different proportions of removed sites, is the ratio of the probability of removing mislabelled sites to the probability of removing non-mislabelled sites. Mathematically, relative risk is defined as:

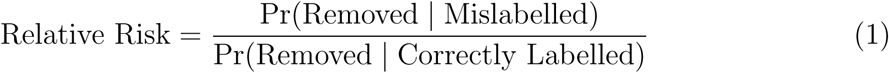

For empirical datasets, given the uncertainty regarding the true tree, we used the RF distance to compare the inferred trees resulting from removing different proportions of sites, to the best-supported trees in the literature. Additionally, we calculated evolutionary rates for each site in the empirical datasets when site-rate heterogeneity models were applied using IQ-TREE2. This is achieved by first estimating model parameters, followed by using an empirical Bayesian approach to assign site-specific rates. The rates are calculated as the weighted average over rate categories, with the weights determined by the posterior probability of each site belonging to a particular category (Minh et al., 2020). For models assuming site-rate homogeneity, we used parsimony scores as a proxy to assess whether the removed sites were fast-evolving.

#### Robinson-Founds distance and branch score

Robinson and Foulds (1981) introduced the RF distance, a metric that compares two phylogenetic trees with the same taxa. This metric measures the differences between two trees by counting the number of splits induced by one of the trees but not the other. An RF distance of zero indicates that the compared trees share identical topologies.

Branch score, a distance measure developed by Kuhner and Felsenstein (1994), sums the squared differences in branch lengths between two trees. If an branch exists in one tree but not the other, the squared difference is the square of the branch length from the tree where the branch is present. The smaller the branch scores, the more similar the branch lengths between the two trees compared.

For the simulation studies, we can determine the optimal cut-off ranges by comparing the inferred trees to the generating trees using the RF distance and the branch score. That is, the cut-off ranges of the percentage of sites removed were identified where the inferred trees show the lowest RF distance and the lowest branch score.

## Results

### Simulation Results

#### Mislabelling simulation results

In the mislabelling simulations, we first evaluate the effectiveness of the trimmed log-likelihood method by investigating whether the sites excluded in the tree inferred were those that had been mislabelled. The relative risks for all three simulations were consistently greater than 1, demonstrating that our trimmed log-likelihood method effectively identified and removed mislabelled sites more frequently than non-mislabelled sites (Fig. 3). Specifically, in the Short branch length simulations, when 1% and 2% of sites were removed, mislabelled sites were three times more as likely to be removed compared to non-mislabelled sites (Fig. 3). As the proportion of sites removed increased, the relative risk decreased to 1.5 at 20%, suggesting that while our method continued to effectively identify mislabelled sites, the distinction between mislabelled and non-mislabelled site removal became less pronounced as the proportion of removed sites increased (Fig. 3).

Similarly, in the Medium branch length simulations, the relative risk were also higher than 3 for both 1% and 2% of sites removed, then decreases to 1.7 when 20% of sites were removed (Fig. 3). In the Long branch length simulations, the relative risk was 2.9 when 1% of sites were removed and then declined to 1.9 by the time 20% of sites were removed (Fig. 3). The lower starting relative risk value in the Long branch length simulations may be explained by the fact that these alignments contained fewer constant sites and more fast-evolving or saturated sites compared to the Short and Medium branch length simulations. The trimmed log-likelihood method targeted sites with the lowest log-likelihood sites, and both erroneous sites and fast-evolving sites tend to have low log-likelihoods. As a result, the likelihood of removing incorrectly labelled sites decreased. This could explain the lower starting relative risk values in the Long branch length simulations compared to the Short and Medium branch length simulations.

In addition, we explored the optimal cut-off points for removing mislabelled sites, as determined by the lowest RF distances to the tree used to simulate the data, which varied across different mislabelling simulations (Fig. 4a). In the Short branch length simulations, the mean RF distances of the inferred tree topologies were lowest when up to 10% of the sites were removed. When the proportion of removed sites increased from 11% to 20%, the mean RF distance increased, as indicated by the error bars in Fig. 4a. In the Medium branch length simulations, the range of the lowest mean RF distance was similar to that of the Short branch length simulations, around 0% to 10% (Fig. 4a). For the Long branch length simulations, the lowest mean RF distance occurred when 10% to 20% of sites were removed (Fig. 4a).

**Figure 4:**
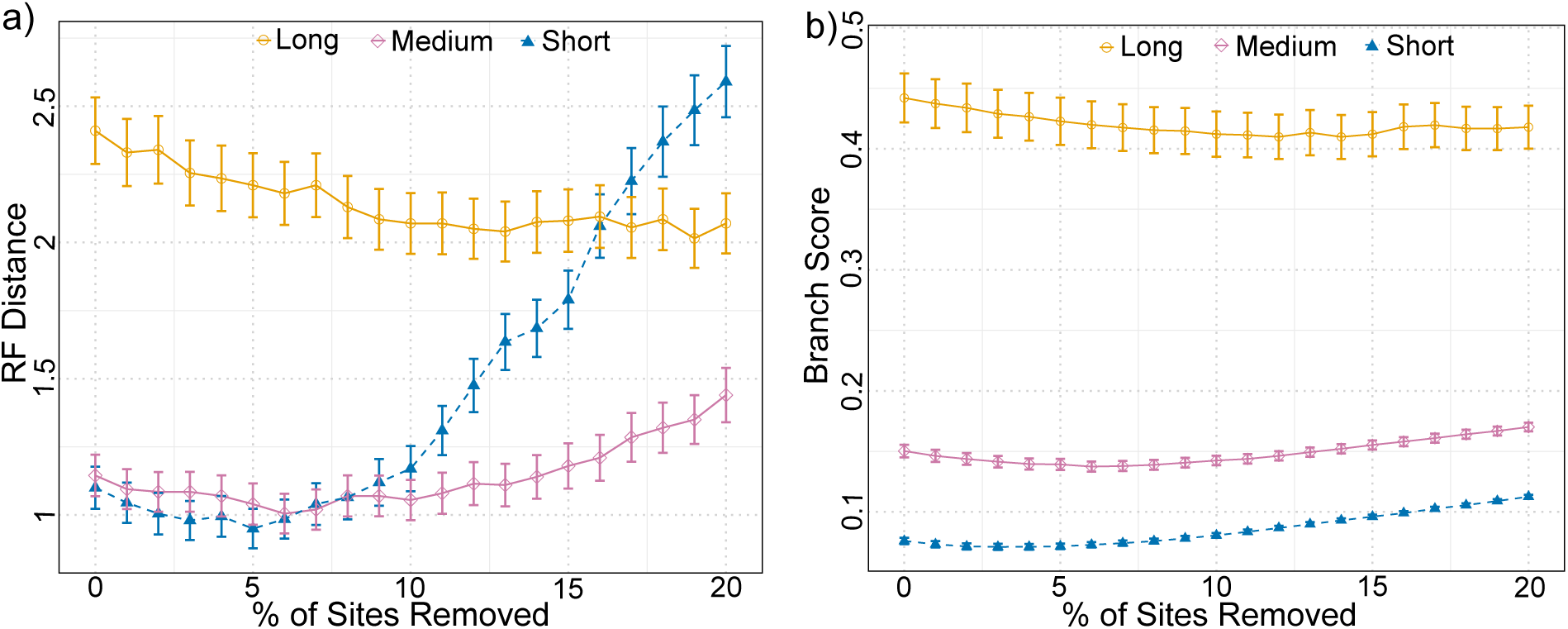
The average Robinson-Foulds (RF) distances and average branch scores between the inferred trees and the generating trees were calculated as varying proportions of sites were removed in each simulation. (a) The mean RF distances for the Long, Medium, and Short branch length simulations. Vertical bars represent *±*1 standard errors of the mean RF distance. (b) The mean branch scores for the three simulations, with vertical bars indicating *±*1 standard errors of the mean branch score.

A similar pattern was observed in the branch scores results. For the Short and Medium branch length simulations, the mean branch scores were lower when removing 0% to 10% of the sites compared to higher removal percentages, such as 15% to 20% (Fig. 4b). In contrast, for the Long branch length simulations, the mean branch scores showed a slight decrease as the proportion of sites removed increased (Fig. 4b).

In summary, these simulations demonstrated the effectiveness of our trimmed loglikelihood method in consistently identifying and removing mislabelled sites across various scenarios. These findings were particularly promising, especially given that some mislabelled sites were constant or singleton. Specifically, in the Short and Medium branch length simulations, the relative risk reached as high as 3.2, underscoring the effectiveness of the approach in targeting and removing mislabelled sites.

The mislabelling simulations showed that as the proportion of fast-evolving sites in the alignment increased, it became less likely for incorrectly labelled sites to be removed. This is because our trimmed log-likelihood method focuses on fast-evolving sites. In the Short and Medium branch length simulations, the relative risk of removing mislabelled sites was higher than in the Long simulations, as these alignments had fewer fast-evolving sites. With more slow-evolving sites like constants and singletons, the method more accurately identified mislabelled sites in these cases.

#### Convergent process simulation results

When applying the trimmed log-likelihood methods and fitting a JC model to the simulated datasets, the accuracy of the inferred tree topologies varied depending on the proportion of sites removed (Fig. 5a-c). For the 0.01-800JC-200GC, 0.01-700JC-300GC, and 0.01-500JC-500GC simulations, the lowest mean RF distances were observed when removing around 0% to 10% of the sites, with higher sites removal leading to increased mean RF distances (Fig. 5a). In the 0.03-800JC-200GC, 0.03-700JC-300GC, and 0.03-500JC-500GC simulations, the lowest mean RF distances occurred with 0% to 15% of the sites removal (Fig. 5b). For the 0.05-800JC-200GC, 0.05-700JC-300GC, and 0.05-500JC-500GC simulations, only the first two had the lowest mean RF distances around 0% to 20% of sites removed, while the last simulation showed minimal difference in mean RF distance across different proportions of sites removal (Fig. 5c).

**Figure 5:**
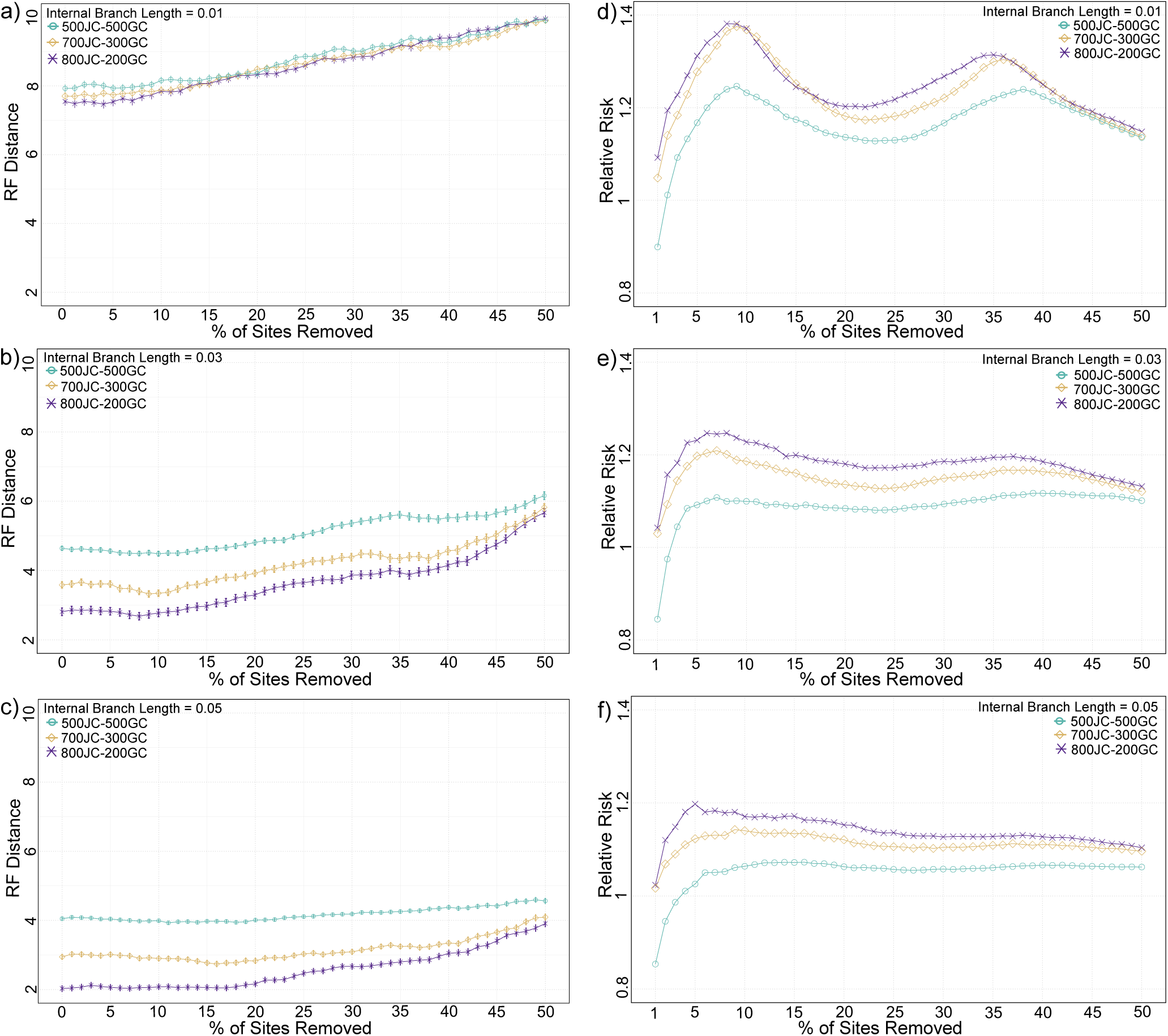
(a-c) Average RF distances for the inferred trees as varying proportions of sites were removed in each simulation. Vertical bars represent *±*1 standard error of the mean RF distance. (d-f) Relative risk of removing GC-biased sites compared to GC-balanced sites at different proportions of sites removed in each simulation.

Across simulations with different internal branch lengths, the trimmed log-likelihood method was most effective in the shortest internal branch length setting (i.e., 0.01), where smaller proportions of sites were removed to achieve the lowest RF distances compared to longer branch length simulations (Fig. 5a-c). Additionally, when comparing different proportions of GC-biased sites, the method was more effective with fewer GC-biased sites in the alignment (i.e., 800JC-200GC), achieving the smallest RF distances with fewer sites removed (Fig. 5a-c). This effect is particularly noticeable in the 0.05-800JC-200GC simulations, where the lowest RF distances occurred with 0% to 20% of sites removed, whereas the 0.05-500JC-500GC showed little change in the RF distances across different levles of sites removal (Fig. 5c).

In contrast to the RF results, branch scores decreased as the proportion of sites removed increased, indicating improved accuracy in the inferred branch lengths (Supplementary Fig. S1). This suggests that the improvement in branch length estimation outweighed the decline in topology accuracy. This is illustrated by two trees inferred from the same alignment in Supplementary Figure S2: Tree 1, inferred using the complete alignment, and Tree 2, inferred after removing 50% of the sites. Compared to the generating tree (Fig. 1), Tree 1 had an RF distance of 4 and a branch score of 1.3, while Tree 2 had an RF distance of 8 but a lower branch score of 0.6. Although Tree 1 had a more accurate topology, Tree 2, with fewer sites, produced more accurate branch lengths (Supplementary Fig. S2).

Furthermore, we also calculated the relative risk of removing GC-biased sites versus GC-balanced sites at different proportions of sites removed. The relative risk was calculated similarly to Equation 1, but by comparing the probability of removing GC-biased sites to the probability of removing GC-balanced sites.

The relative risks for the 800JC-200GC and 700JC-300 GC simulations were always greater than 1 but less than 1.4, suggesting that our trimmed log-likelihood method had limited effectiveness in these cases, particularly compared to the mislabelling simulations (Fig. 5d-f). In the 500JC-500GC simulations, removing around 1% to 2% of sites, the relative risk was below 1, showing that the method was ineffective in these instances (Fig. 5d-f).

The relative risk results indicated the limited effectiveness of the trimmed log-likelihood method in recovering true tree topologies, mainly due to its inability to effectively remove GC-biased sites. The relative risk showed only a small increase when a small proportion of sites was removed - around 1% to 10% in the 800JC-200GC and 700JC-300GC simulations, or around 3% to 10% in the 500JC-500GC simulations. The marginal increase suggested that GC-biased sites were only slightly more likely to be removed than GC-balanced sites. As a result, the remaining GC-biased sites contributed to incorrect tree topologies.

#### Saturation simulation results

Our simulation results showed that removing from 18% to 30% of the sites led to the lowest average RF distance between the inferred trees and the generating tree (Fig. 6b). Specifically, removing sites within the intervals 19% to 20%, 23% to 28%, and 47% to 50% resulted in consensus trees with an RF distance of 2 compared to the generating tree (Fig. 6a). This indicates that these consensus trees successfully recovered all splits except for the split “ABCD|EFG” (Fig. 2a). These results are similar to those reported by Cummins and McInerney (2011), who found that using the TIGER approach and removing between 183 and 502 fast-evolving sites (approximately 18% to 50% of total sites) resulted in consensus trees that recovered all splits in the generating tree except for the “ABCD|EFG” split. Our methods demonstrated effectiveness comparable to that of the TIGER approach.

**Figure 6:**
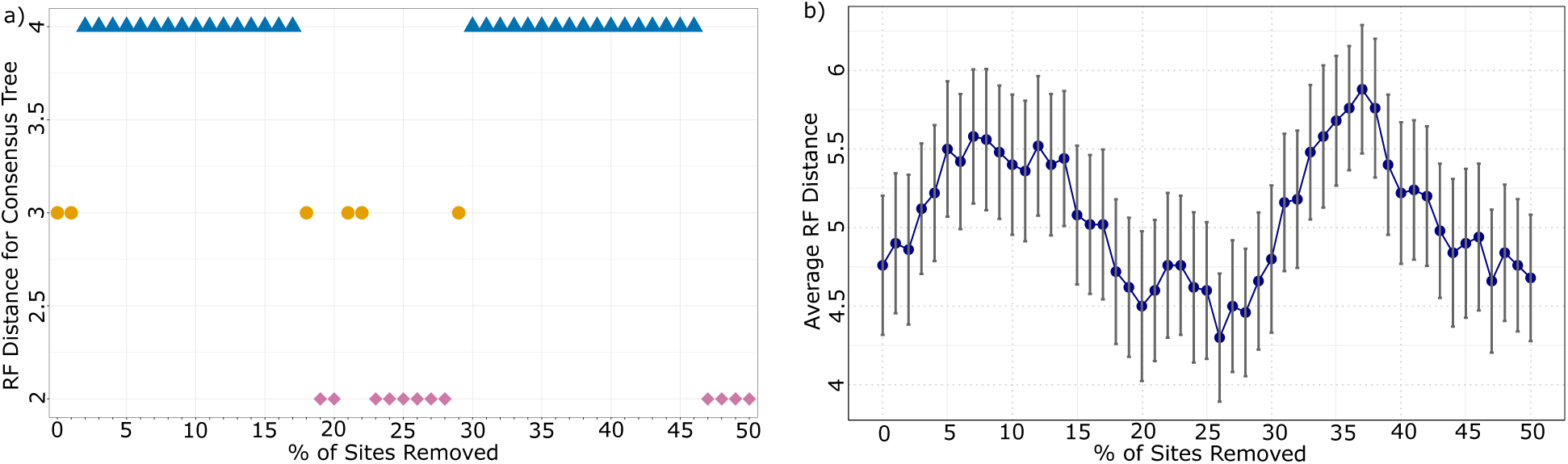
Saturation simulation results. (a) The RF distance between the generating tree and the consensus tree constructing from the 100 inferred trees for each proportion of sites removed. The diamond shaped points indicate cases where the consensus tree has a RF distance of 2, meaning it recovered the generating tree topology except the “ABCD|EFG” split. (b) Average RF distance between the generating tree and the inferred trees for different proportion of sites removed.

However, for certain intermediate proportions, such as 21% to 22%, and 29% to 46%, the consensus trees had an RF distance of 3 or 4, failing to recover most phylogenetic relationships in the generating tree. One possible explanation is that the sites removed in these ranges included informative sites, those that strongly supported the generating tree topology but had relatively lower log-likelihood values. The trimmed log-likelihood method, while effective at removing low-likelihood sites, may have mistakenly removed these informative sites, leading to a less accurate inference of the tree.

In summary, the results showed that our method was ineffective in this saturation simulation because it required removing a large number of sites to achieve the best consensus trees, which recovered all but one split of the generating tree topology. This limitation was due to the presence of large amounts of saturated sites in the alignments. As expected, the trimmed log-likelihood method, designed to identify sites with the lowest log-likelihood, struggled to differentiate between saturated sites, all of which had very low log-likelihood values. Similarly, the TIGER approach from Cummins and McInerney (2011) faced the same challenges with alignments containing mostly saturated sites, also requiring the removal of many sites to construct consensus trees that nearly matched the generating tree topology, missing only one split.

### Empirical Dataset Analysis Results

#### Thermus dataset analysis

Using the trimmed log-likelihood method, we applied three different substitution models, TIM2+F+G4, GTR+G+I, and JC, to the *Thermus* dataset. When using the complete dataset without trimming any sites, the RF distance between the inferred tree and the Reference tree is 2 (Fig. 7a-c). In fact, with this complete dataset, the topology of the inferred tree was the same as the topology of the Attract tree, grouping the two GC-rich taxa, *Deinococcus* and *Bacillus*, together (Supplementary Fig. S3b). Removing 1% to 2% of sites and applying the TIM2+F+G4 model, we achieved an RF distance of 0 between the inferred tree and the Reference tree is 0, successfully reconstructed the Reference tree (Fig. 7a). Similarly, using the GTR+G+I model and removing 2% of sites, the RF distance was also 0, leading to the recovery of the Reference tree (Fig. 7b). Finally, the removal of 4% to 9% of sites using the JC model also led to the successful reconstruction of the Reference tree (Fig. 7c). It is worth noting that, regardless of the model applied in the trimmed log-likelihood method, the topologies of the inferred trees with the same RF distance were identical (Supplementary Fig. S3).

**Figure 7:**
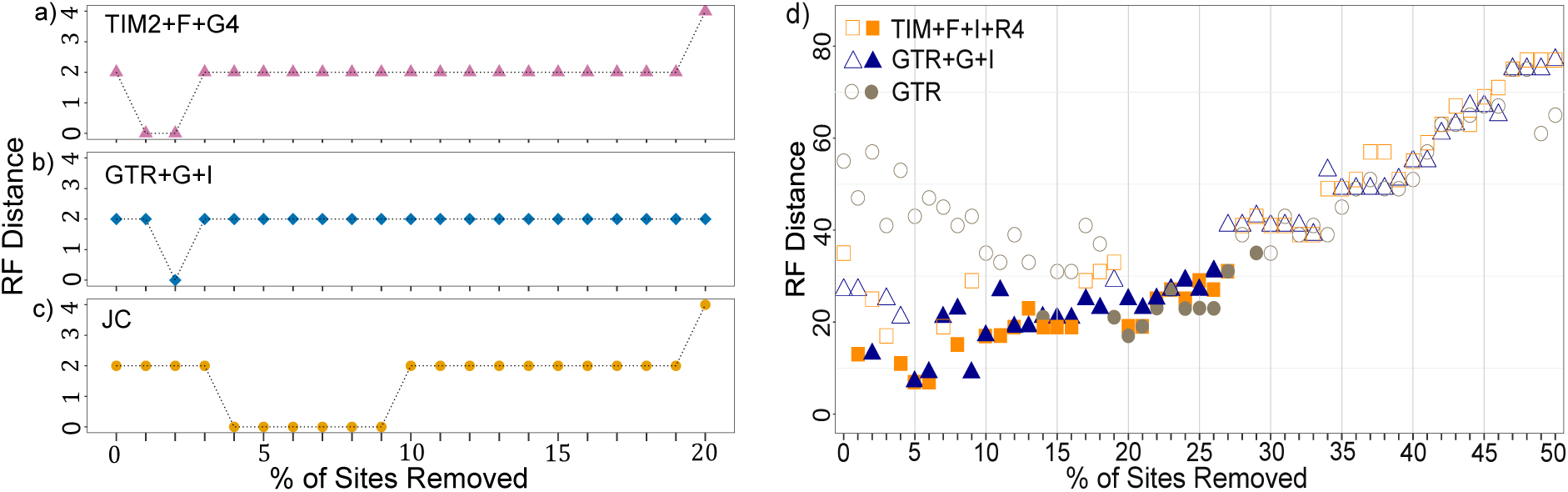
(a-c) RF distances between the trees inferred from the *Thermus* dataset using the (a) TIM2+F+G4, (b) GTR+G+I and (c) JC models, compared to the Reference tree topology (which grouped *Deinococcus radiodurans* and *Thermus aquaticus* as a sister group). Note that all inferred trees with an RF distances of 2 had the same topology as the Attract tree topology, which grouped *Deinococcus radiodurans* and *Bacillus subtilis* as a sister group. (d) RF distances between the trees inferred from the arthropod dataset using the TIM+F+I+R4, GTR+G+I and GTR models, compared to the strict consensus tree shown in Figure 1 in Pisani (2004). Solid dots indicate that the inferred trees support the *Mandibulata* hypothesis.

In comparison, Cummins and McInerney (2011) used the TIGER approach, which involved removing 108 fast-evolving sites (approximately 8% of the total). The GTR+G+I model was applied to the remaining data, recovering the Reference tree. Our trimmed log-likelihood method proved more efficient, requiring only the removal of 2% of sites, focusing on the most relevant ones, to recover the correct topology using the same model (Fig. 7b). Even with the simplest substitution model, i.e., the JC model, the Reference topology could be recovered by removing just 4% of the lowest-likelihood sites (Fig. 7c). This analysis showed that the better the model, as determined by AIC values, required fewer sites to be removed. For example, the TIM2+F+G4 model, which was the best-fit model according to AIC, required the removal of only 1% - 2% of sites (Fig. 7a). In contrast, the JC model required the removal of 4% - 9% (Fig. 7c).

We further investigated the nature of the sites removed by the trimmed log-likelihood method to determine whether they were fast-evolving sites. We calculated the evolutionary rates of the removed sites when applying the TIM2+F+G4 and GTR+G+I models to the *Thermus* dataset in IQ-TREE2. Additionally, for the JC model, parsimony scores were used as a proxy for evolutionary rates, as that model does not allow for rate categories to be assigned to sites. Specifically, we examined the 1% of sites removed using the TIM2+F+G4 model, the 2% of sites removed using the GTR+G+I model, and the 4% of sites removed using the JC model (Fig. 8a-c). In all cases, the removed sites had high evolutionary rates, confirming that they were fast-evolving sites (Fig. 8a-c).

**Figure 8:**
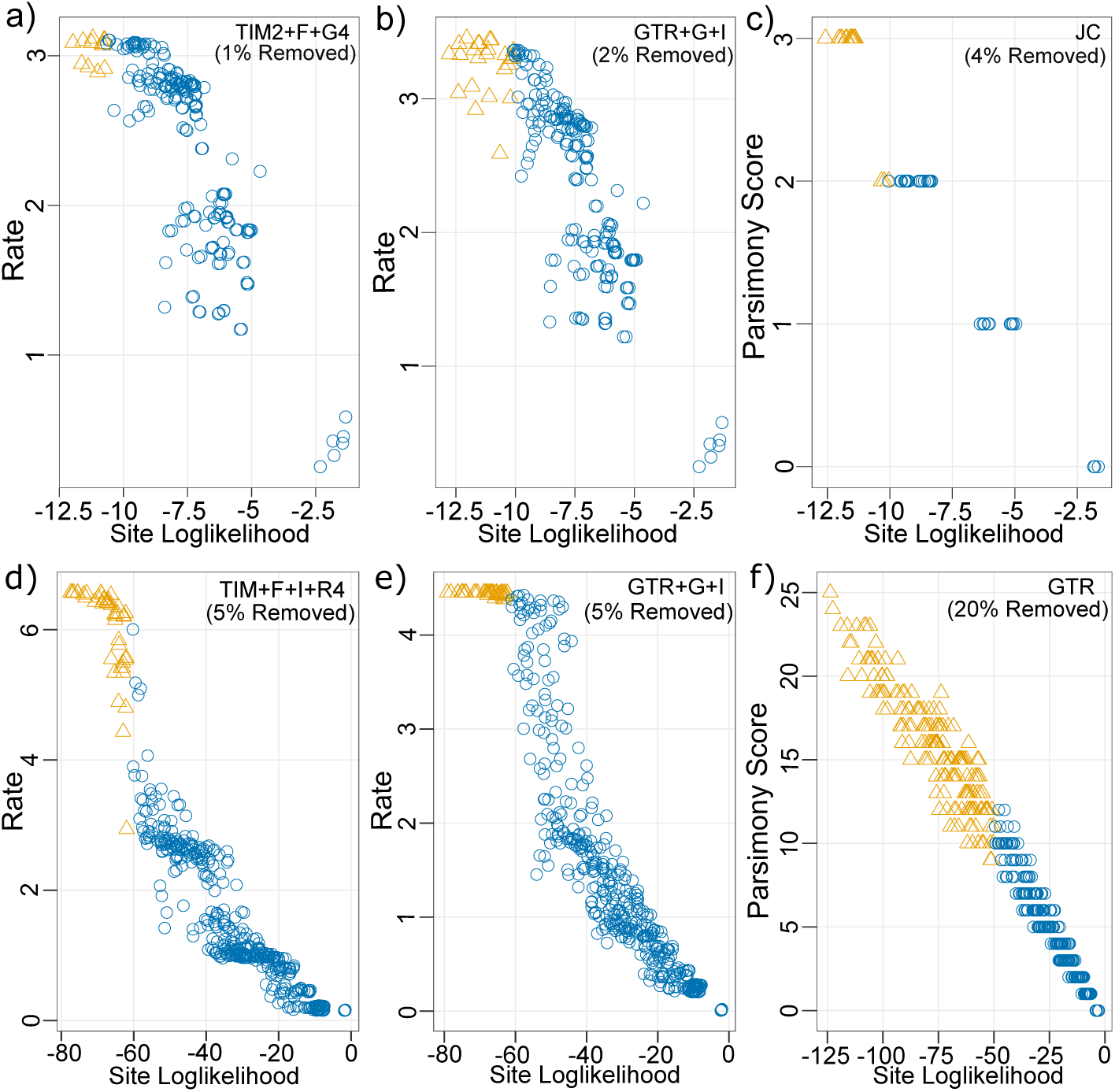
Site evolutionary rates and log-likelihoods for three different models in the *Thermus* dataset (a-c) and arthropod dataset (d-f) analyses. (a-b, d-e) Site-specific evolutionary rates were calculated using IQ-TREE2 with the TIM2+F+G4 and GTR+G+I models for the *Thermus* dataset, and the TIM2+F+I+R4 and GTR+G+I models for the Arthropod dataset. (c, f) Parsimony scores, used as approximations for evolutionary rates, were applied with the JC model in the *Thermus* dataset and the GTR model in the arthropod dataset, both with varying proportions of sites removed. Triangular dots indicate the removed sites.

Moreover, given that the three thermophiles (*Aquifex arolicus*, *Thermatoga martima* and *Thermus aquaticus*) in the *Thermus* dataset had higher GC content than the two mesophiles, we aimed to examine whether the sites removed during the analysis were also biased the towards GC-rich region. For the 1% of sites (12 sites) removed under the TIM2+F+G4 model, it was unclear that the removed sites were biased toward GC-rich regions in these three thermophiles (Supplementary Fig. S3d). This is likely due to the small number of sites removed, making it difficult to identify any clear pattern. However, with further inspection of the specific sites removed using the TIM2+F+G4 model, we found that these excluded sites supported the grouping of the two mesophiles, *Bacillus subtilis* and *Deinococcus radiodurans*, where the taxa either had matching nucleotides or were in the same purine/pyrimidine class, reflecting their more balanced GC content (Supplementary Table S1).

In contrast, when removing 2% and 4% of sites (25 and 50 sites, respectively) under the GTR+G+I and JC models, we observed a clearer pattern: the removed sites consistently showed higher GC content in the thermophiles compared to the mesophiles (Supplementary Fig. S3e and S3f). For example, in the complete alignment, the three thermophiles have approximately 28% and 36% for C and G nucleotides, respectively. However, the sites removed using the JC model showed equal or even higher GC content - ranging from 34% to 44% for C nucleotides and 32% to 38% for G nucleotides (Supplementary Fig. S3f).

In summary, the trimmed log-likelihood method, applied across different substitution models, demonstrated its effectiveness in recovering the reference phylogeny by removing small percentages of fast-evolving sites. Importantly, the removed sites also included regions associated with the three thermophilic taxa, which are rich in GC content. The trimmed log-likelihood method not only successfully recovered the Reference tree but also performed more efficiently than methods like TIGER by requiring the removal of fewer sites while achieving comparable accuracy.

#### Arthropod dataset analysis results

By applying the trimmed log-likelihood method and fitting three different models to the arthropod dataset, we successfully recovered the relationship among the *Chelicerata*, *Myriapoda* and *Pancrustacea* clades, supporting for the *Mandibulata* hypothesis (Fig. 7d and Supplementary Fig. S6, S8 and S10). For the best-fit model chosen by AIC (TIM+F+I+R4), the lowest RF distance (RF = 7) was achieved by removing 5% and 6% of sites (43 and 52 sites) (Fig. 7d). Similarly, under the GTR+G+I model, the same lowest RF distance (RF = 7) was obtained after removing 5% of sites. These optimal inferred trees, despite having an RF distance of 7, closely resembled the strict consensus tree after removing 5% or 6% of sites. The differences between the optimal trees (Supplementary Fig. S6 and S8) and the strict consensus tree (Supplementary Fig. S4) came from ambiguous relationships observed in some external branches of the strict consensus tree, such as *Polycentropus interruptus*, *Byrsopteryx gomezi*, and *Opthalmopsylla volgensis*.

Furthermore, removing other proportions of sites using the TIM+F+I+R4 and GTR+G+I models also successfully recovered the *Mandibulata* hypothesis (Fig. 7d). For TIM+F+I+R4, trees inferred after removing 1%, 4% - 6%, 8%, 10%- 16%, 20% - 27%, recovered the *Mandibulata* hypothesis (Fig. 7d). For the GTR+G+I models, the *Mandibulata* hypothesis was recovered after removing 2%, 5% - 18%, 20% -26% of sites (Fig. 7d).

We also applied a simpler and poorer-fitting model, GTR, to the arthropod dataset. After removing 20% of sites (173 sites), the inferred tree achieved the lowest RF distance of 17 (Fig. 7d). Despite using this poorer-fitting GTR model, the trimmed log-likelihood approach successfully recovered the relationships among the *Chelicerata*, *Myriapoda* and *Pancrustacea* clades relationship, thereby supporting the *Mandibulata* hypothesis (Supplementary Fig. S10). Additionally, removing 14%, 19% -27% and 29% also led to recovery of the *Mandibulata* hypothesis (Fig. 7d).

When analysing the tree inferred using all sites with these three models, we observed discrepancies in the recovery of the relationships among the *Chelicerata*, *Myriapoda* and *Pancrustacea* clades. The TIM+F+I+R4 and the GTR+G+I models misplaced two *Hexapoda* species, *Ctenolepisma lineata* and *Litobius forficatus*, within the *Myriapoda* clade (Supplementary Fig. S5 and S7). The GTR model performed even worse when using the entire dataset: it incorrectly grouped *Myriapodas* and *Chelicerates* together, which supports the *Myriochelata* hypothesis. Furthermore, some species from the *Pancrustacea* clade were incorrectly placed within either the *Myriapoda* or *Chelicerate* clade (Supplementary Fig. S9).

In comparison to the results presented in Pisani (2004), where the authors used the SF method (Brinkmann and Philippe, 1999) and the LQP value (Wilkinson, 1992) to identify fast-evolving sites in the Neighbour-Joining (NJ) analysis, their studies found that removing sites based on the LQP value helped to improve the accuracy of the inferred trees in the NJ analysis. The NJ tree, inferred after excluding 172 sites (*≈* 20%) with an LQP *≥* 0.5 (referred to as the best LQP tree), although recovering the *Mandibulata* hypothesis, still had an RF distance of 31 when compared to the strict consensus tree.

Similarly, the optimal NJ tree, after removing 196 sites (*≈* 23%) identified by the SF method (referred to as the best SF tree), also showed an RF distance of 31 (Pisani, 2004). In other words, our trimmed log-likelihood method proved more effective in removing fast-evolving sites for several reasons: firstly, we only needed to remove 43 sites (5%) from the data to obtain a tree supporting the *Mandibulata* hypothesis (Fig. 7d). Secondly, the best inferred trees using three different models had much smaller RF distances (RF = 7 or 17) than the optimal LQP and SF trees in Pisani (2004) (Fig. 7d). Lastly, even when using a poorer-fitting model (i.e., GTR) for the data, our method consistently recovered the true relationships among the three clades mentioned in the *Mandibulata* hypothesis (Supplementary Fig. S10).

Additionally, we investigated whether the sites removed were fast-evolving. Specifically, for the 5% of sites removed using the TIM+F+I+R4 model, the 5% of sites removed using the GTR+G+I model, and the 20% of sites removed using the GTR model, we calculated their site-specific evolutionary rates for the first two models in IQ-TREE2 and used parsimony scores as a proxy for the site-specific evolutionary rate in the GTR model, as that model does not allow us to assign rate categories to sites. We found that the sites removed all had very high evolutionary rates, confirming that they are indeed fast-evolving sites (Fig. 8d-f).

In summary, the trimmed log-likelihood method applied to the arthropod dataset successfully supported the *Mandibulata* hypothesis by recovering the relationships among the *Chelicerata*, *Myriapoda*, and *Pancrustacea* clades. This method outperformed the SF and LQP approaches from Pisani (2004), requiring fewer sites to be removed (as little as 5%) to achieve a tree with lower RF distances (RF = 7 or 17) compared to the optimal NJ trees. Even when using poorer-fitting models, like GTR, the trimmed log-likelihood method consistently supported the *Mandibulata* hypothesis after removing 20% of sites. Furthermore, the sites removed were confirmed to be fast-evolving, further validating the effectiveness of the trimmed log-likelihood method.

## Discussion

Our simulation studies and empirical dataset analyses have demonstrated that our trimmed log-likelihood method effectively identifies fast-evolving or erroneous sites (e.g. mislabelling sites) (Fig. 5 and 8). However, determining the optimal number of sites to remove to infer the best phylogenies remains unresolved (e.g., Fig. 7). Various factors can make certain sites misleading for phylogenetic inference, and since the true tree is unknown, it is challenging to establish a threshold for site removal.

In addition, our mislabelling simulations, convergent process simulations, and saturation simulations suggest a pattern: the longer the branch lengths of the underlying trees, the larger the proportion of sites that need to be removed to improve inferred topologies (Fig. 4, 5 and 6b). In particular, this pattern is shown in the mislabelling simulation results, where the lowest mean RF distances in the Long branch length simulations were achieved by removing a larger proportion of sites compared to the Short and Medium branch length simulations (Fig. 4).

This challenge of determining how many sites to remove is not unique to our method. The same issue arises when using methods like TIGER, SF and LQP methods. Similar to these approaches, we recommend comparing inferred trees from alignments with varying numbers of removed sites when using the trimmed log-likelihood method. It is particularly important to monitor substantial changes in tree topology resulting from site removal, as these changes may indicate either more accurate or misleading tree configurations. In empirical datasets, assessing the plausibility of these changes in inferred trees after site removal requires drawing on existing knowledge of well-supported phylogenetic relationships from the literature.

Rather than focusing solely on how many sites to be removed, it is important to identify which sites to remove. Studies have shown the importance of sites selection in phylogenetic inference (Shavit Grievink et al., 2013; Shen et al., 2017; Francis and Canfield, 2020). For instance, both Shen et al. (2017) and Francis and Canfield (2020) found that removing even a small subset of specific sites from empirical multi-gene alignments can significantly alter phylogenetic outcomes. Our trimmed log-likelihood method focuses on identifying which sites to remove, rather than specifying the number to exclude. As demonstrated by both our simulation results and empirical analyses, our method effectively removed the erroneous sites in the mislabelling simulations and fast-evolving sites in the two empirical datasets. In particular, our method performed well even when applied to alignments like the *Thermus* dataset, which included two taxa with high GC content.

An interesting observation from the *Thermus* dataset is that, while our method struggled to identify GC-biased taxa in the convergent process simulations, it effectively removed GC-rich sites in the three thermophilic taxa (Supplementary S3d-f). This outcome could be attributed to the fact that the removed sites in the *Thermus* dataset were fast-evolving and also tended to be GC-rich, particularly in the thermophilic taxa.

In some of the convergent process simulations, the trimmed log-likelihood method led to only minor changes in the mean RF distance when removing a small proportion of sites (Fig. 5a-c). However, the approach made a significant improvement in the inferred branch lengths (Supplementary Fig. S1). This suggests that even with minimal changes in the tree topology, the trimmed log-likelihood method can still improve the accuracy of certain tree features, such as branch lengths.

Our method offers three distinct advantages over existing approaches. Firstly, it has demonstrated comparable or better performance in identifying fast-evolving sites compared to methods such as the SF, TIGER, and LQP (Fig. 6, 7 and 8). These advantages are reflected in the smaller number of sites removed by our method while still inferring optimal trees (Fig. 7). Secondly, our method avoids circularity by dynamically adjusting the subset of removed sites throughout the estimation of tree topology, model of evolution, and branch lengths. Rather than selecting sites removal based on an initial tree, the method continuously updates site removal throughout the process. Lastly, our method is implemented in IQ-TREE 2, allowing for easy use through a single command line and offering a user-friendly experience compared to other methods.

It is important to note that, despite its strengths, our method faces challenges in identifying sites that contribute to heterogeneous evolutionary processes, particularly with regard to compositional biases in the alignments, such as GC bias (Fig. 5). This is likely to be a general result, because the method removes sites with the lowest likelihoods from each iteration of the tree search. In practice, the largest factor determining the likelihood of a site is usually the number of substitutions required to explain a site on the current tree and given the current model, and is usually rather less affected by the relative probability of each of these substitutions as determined by the model of sequence evolution. Therefore, the method will often remove fast-evolving sites, even if these are not erroneous, in preference to sites that violate the model but evolve more slowly.

Another limitation of the trimmed log-likelihood approach is that it removes entire columns of alignments, making it unable to detect taxon-specific sites — those that impact only certain taxa rather than the entire alignment. This limitation can reduce the effectiveness of the analysis in GC-rich regions and other compositional biased contexts. One potential solution to address this issue could involve removing rogue taxa, which may mitigate the impact of these problematic sites.

Despite these limitations, our method remains valuable for addressing specific challenges in phylogenetic inference. This approach, alongside careful consideration and application of existing knowledge, enhances trimmed log-likelihood analyses across diverse biological contexts. Additionally, the method has potential to serve as a residual diagnostic tool. For instance, plotting the locations of down-weighted sites and identifying clusters could reveal sections of the sequence alignment that merit further attention.

## Data Availability

The datasets, R code, Bash scripts, and supplementary figures and tables used in this study are publicly available at this GitHub repository.

## Supplementary Materials

**Table S1:**
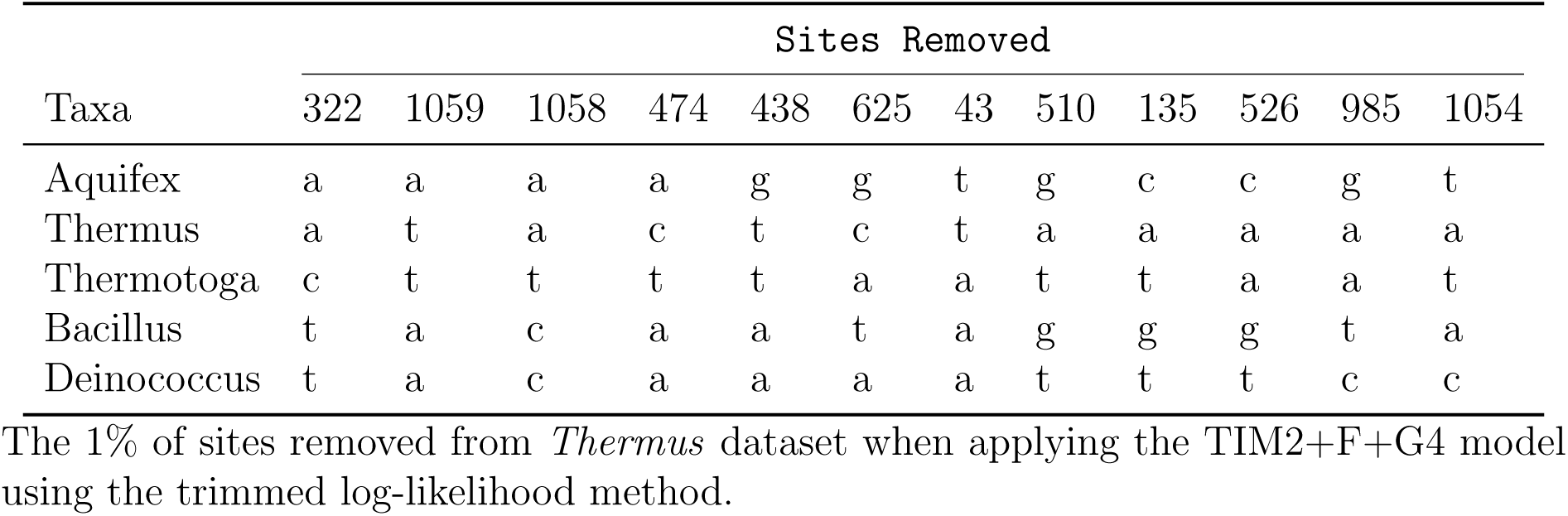
The 1% Sites Removed with TIM2+F+G4 Model.

| Taxa | Sites Removed |  |  |  |  |  |  |  |  |  |  |  |
| --- | --- | --- | --- | --- | --- | --- | --- | --- | --- | --- | --- | --- |
|  | 322 | 1059 | 1058 | 474 | 438 | 625 | 43 | 510 | 135 | 526 | 985 | 1054 |
| Aquifex | a | a | a | a | g | g | t | g | c | c | g | t |
| Thermus | a | t | a | c | t | c | t | a | a | a | a | a |
| Thermotoga | c | t | t | t | t | a | a | t | t | a | a | t |
| Bacillus | t | a | c | a | a | t | a | g | g | g | t | a |
| Deinococcus | t | a | c | a | a | a | a | t | t | t | c | c |

The 1% of sites removed from *Thermus* dataset when applying the TIM2+F+G4 model using the trimmed log-likelihood method.

**Supplementary Figure S1:**
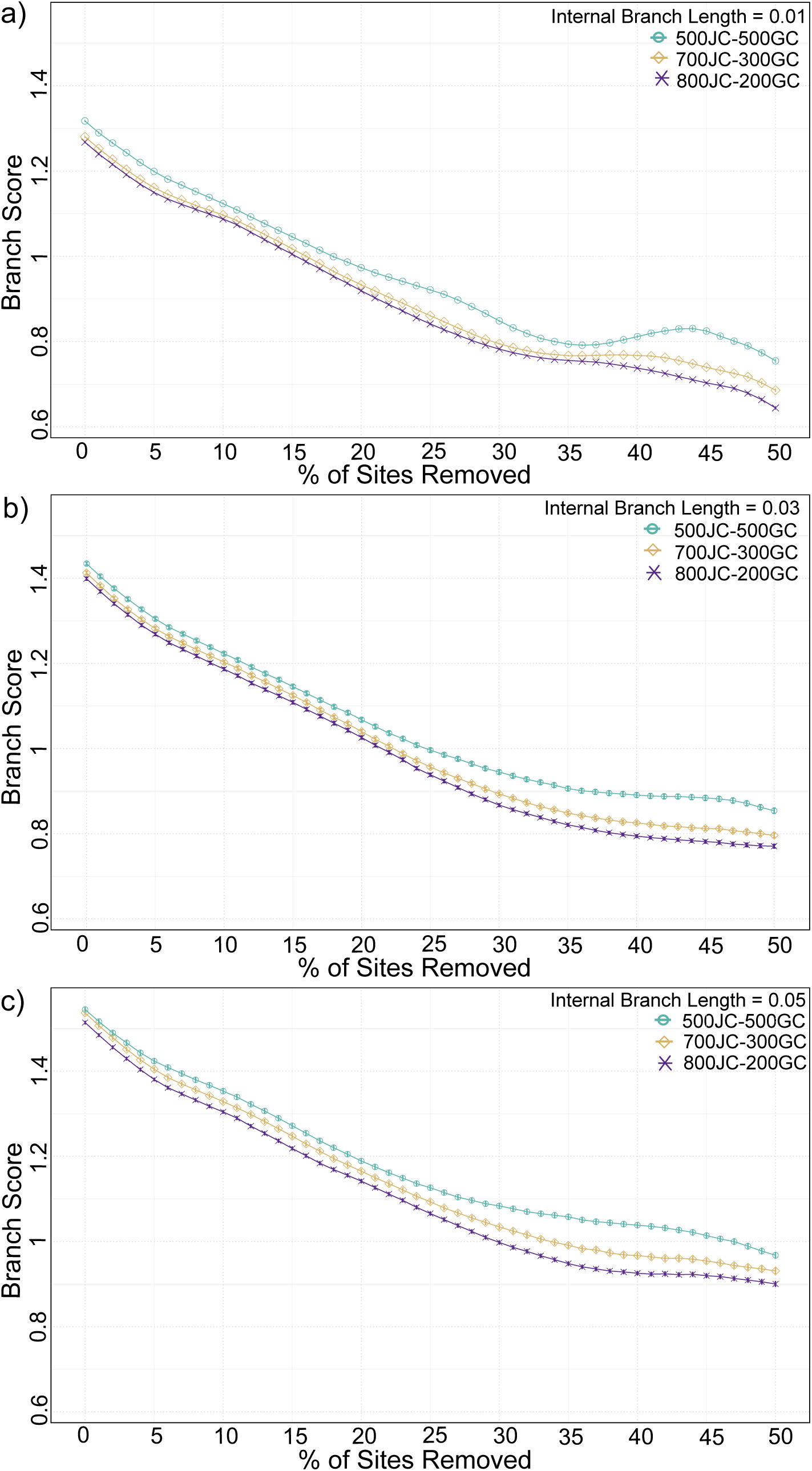
The mean branch scores for the inferred trees as varying proportions of sites were removed in each simulation, with vertical bars indicating *±*1 standard error. Note that standard errors were relatively small compared to the branch scores.

**Figure S2:**
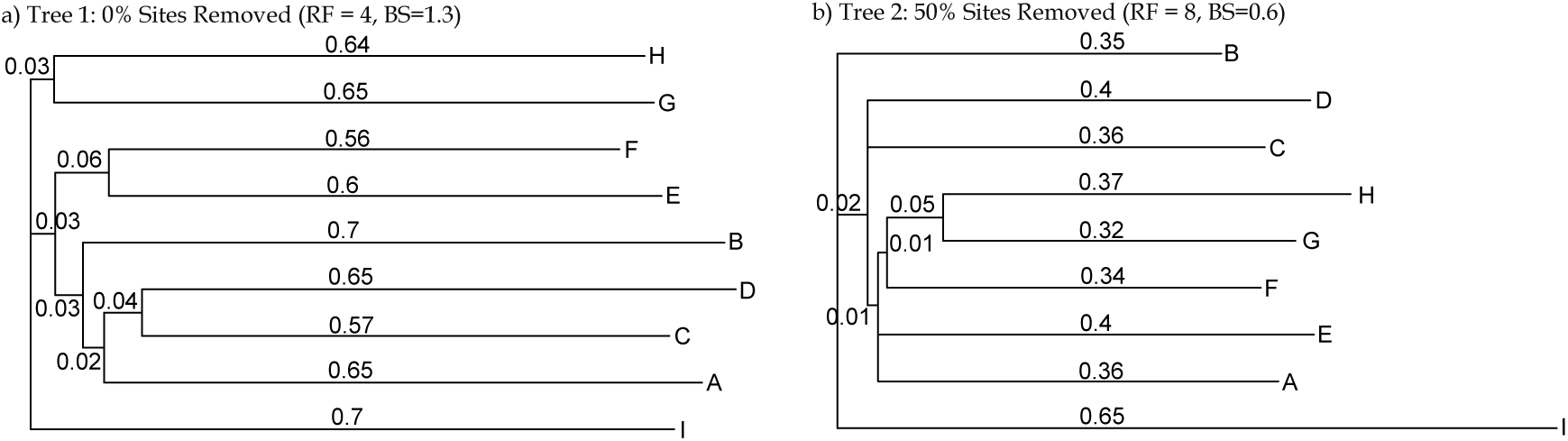
Trees inferred from a simulated alignment: (a) Tree 1 was inferred from the complete alignment in the 0.01-800JC-200GC simulation, with an RF distance of 4 and a branch score of 1.3 compared to the generating tree. (b) Tree 2 was inferred from the same alignment after removing 50% of the sites, with an RF distance of 8 and a branch score of 0.6 compared to the generating tree.

**Figure S3:**
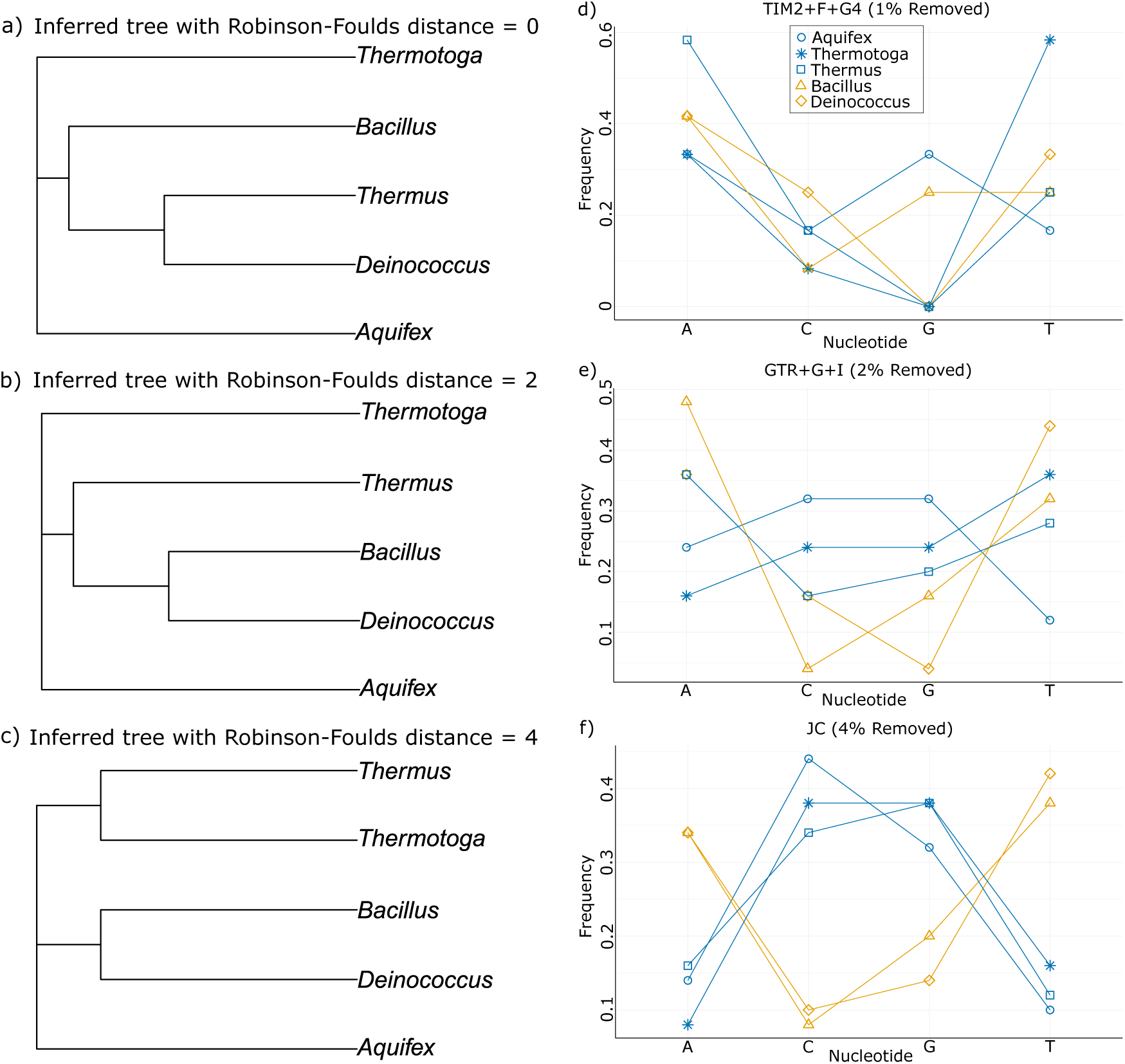
(a–c) Trees inferred from the *Thermus* dataset. The topologies of the inferred trees were identical for each RF distance, regardless of the model used. The figure shows trees with RF distances of 0, 2, and 4, respectively. (d-f) Nucleotide frequencies for 1%, 2%, and 4% of sites were removed using the TIM2+F+G4, GTR+G+I and JC models, respectively, in the *Thermus* dataset.

**Figure S4:**
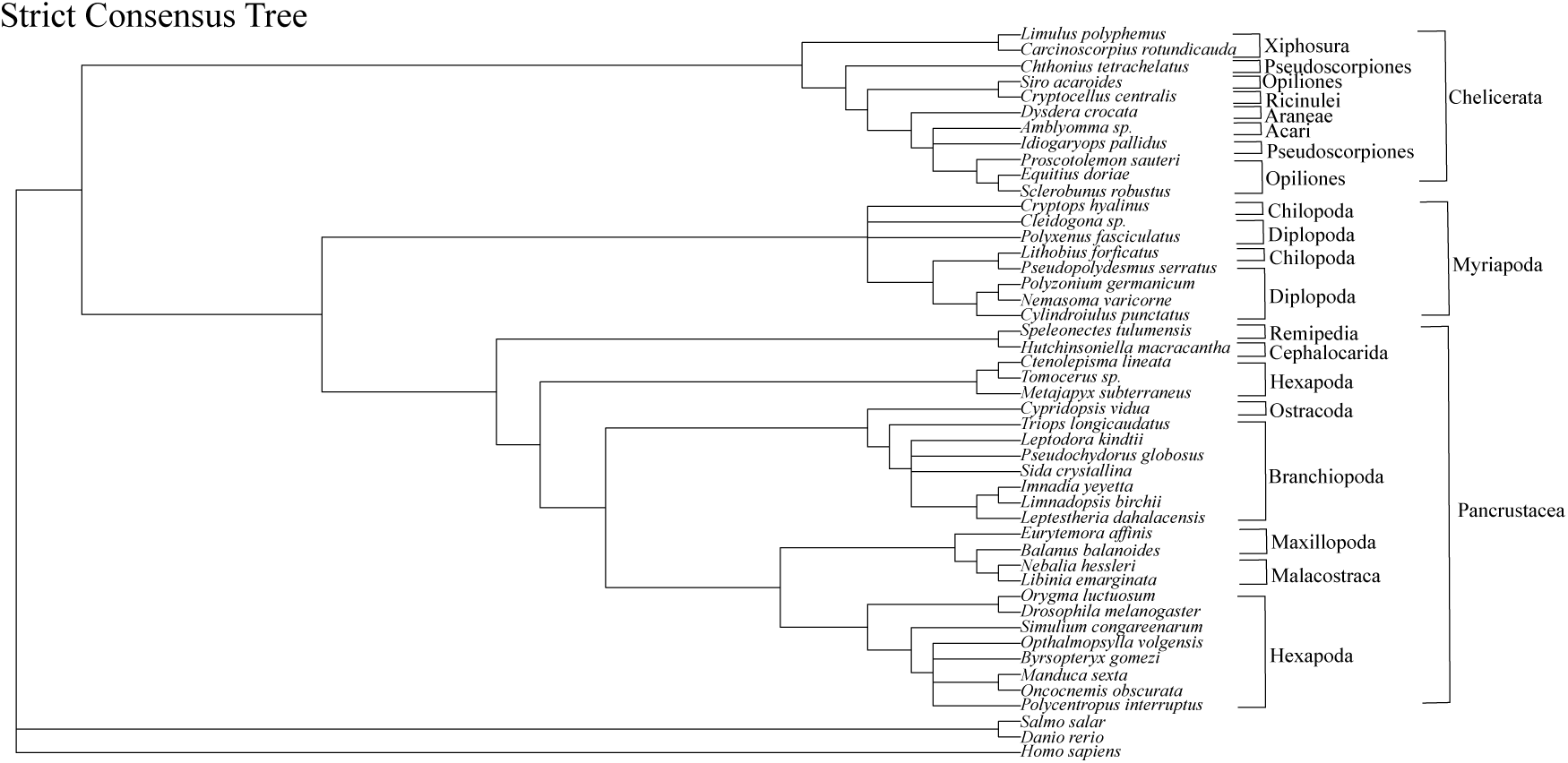
Strict consensus tree from Figure 1 in Pisani (2004), used as a reference for comparison with trees inferred using the TIM+F+I+R4, GTR+ G+I, and GTR models from the arthropod dataset.

**Figure S5:**
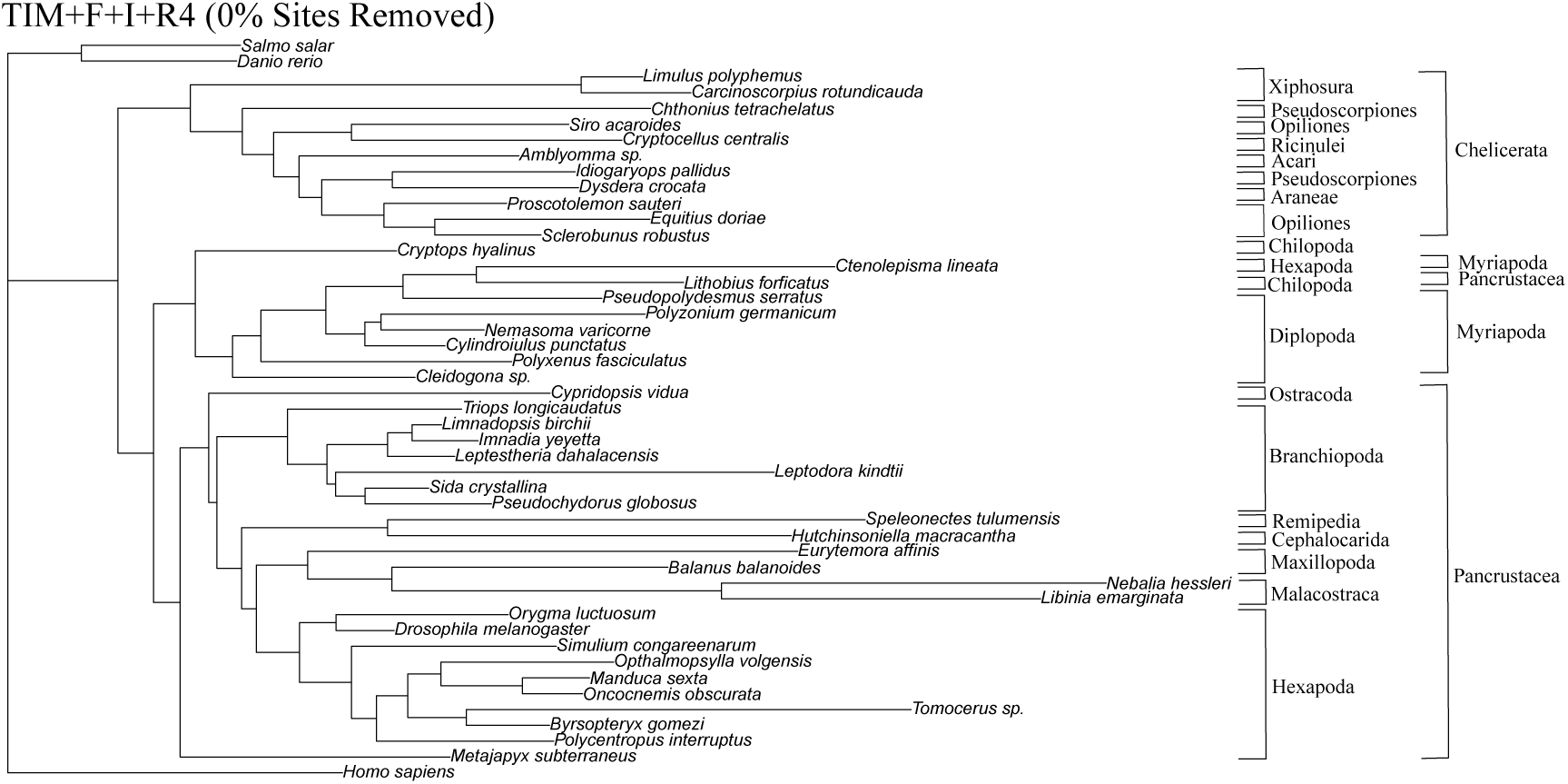
The tree inferred using the TIM+F+I+R4 model on the complete arthropod dataset.

**Figure S6:**
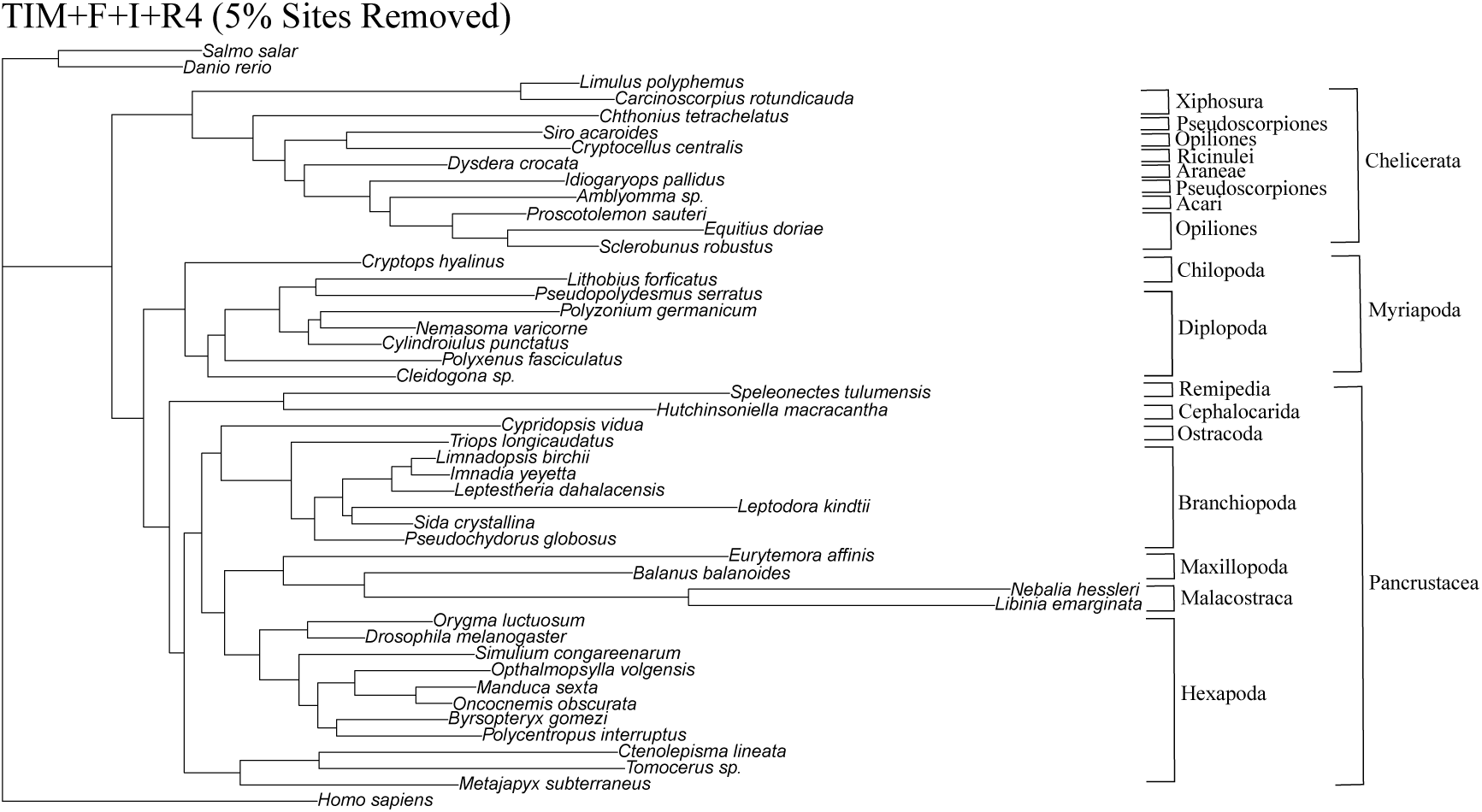
The tree inferred using the TIM+F+I+R4 model after removing 5% of the sites from the arthropod dataset. This is the optimal tree obtained with the TIM+F+I+R4 model.

**Figure S7:**
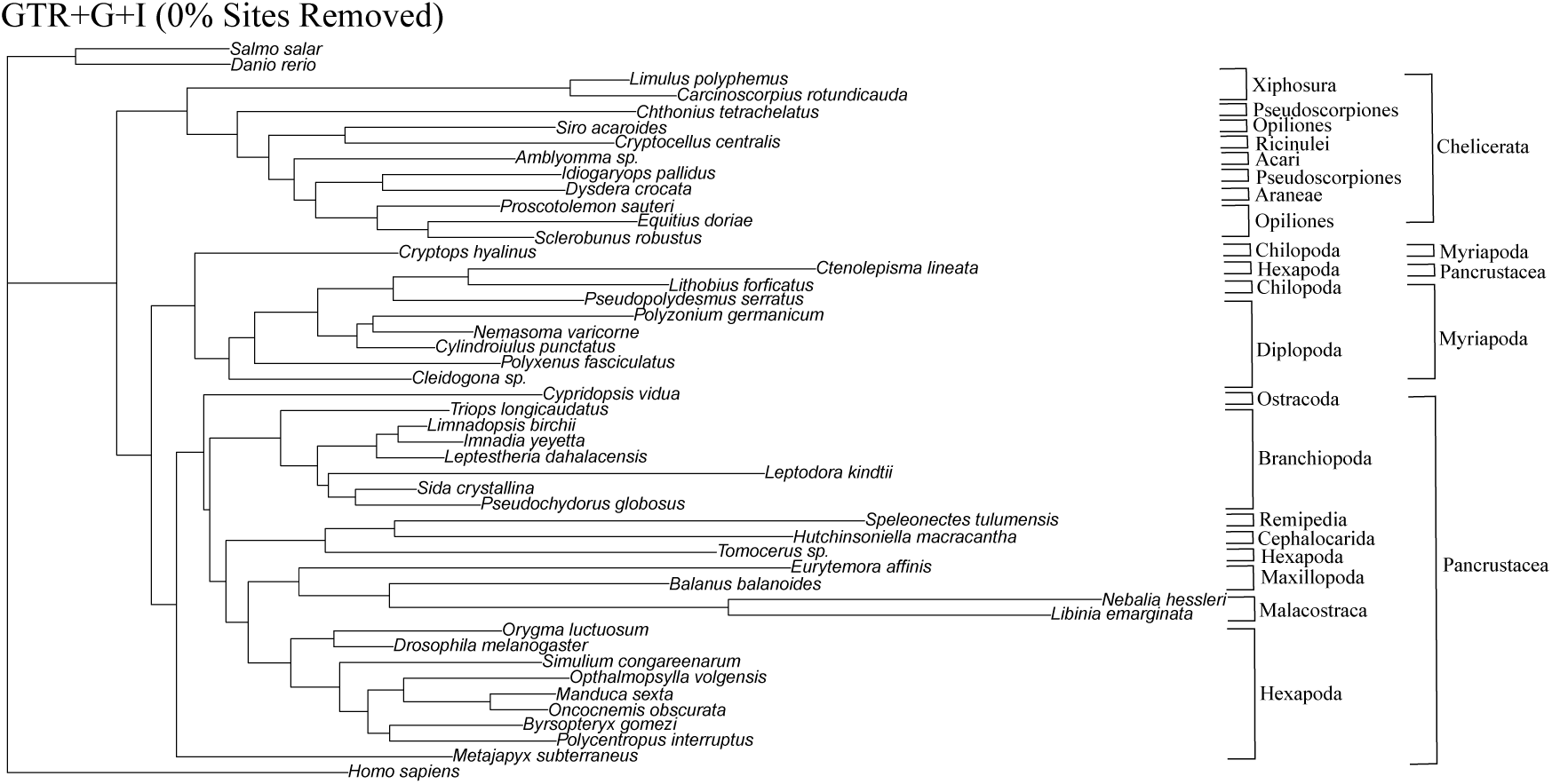
The tree inferred using the GTR+G+I model on the complete dataset.

**Figure S8:**
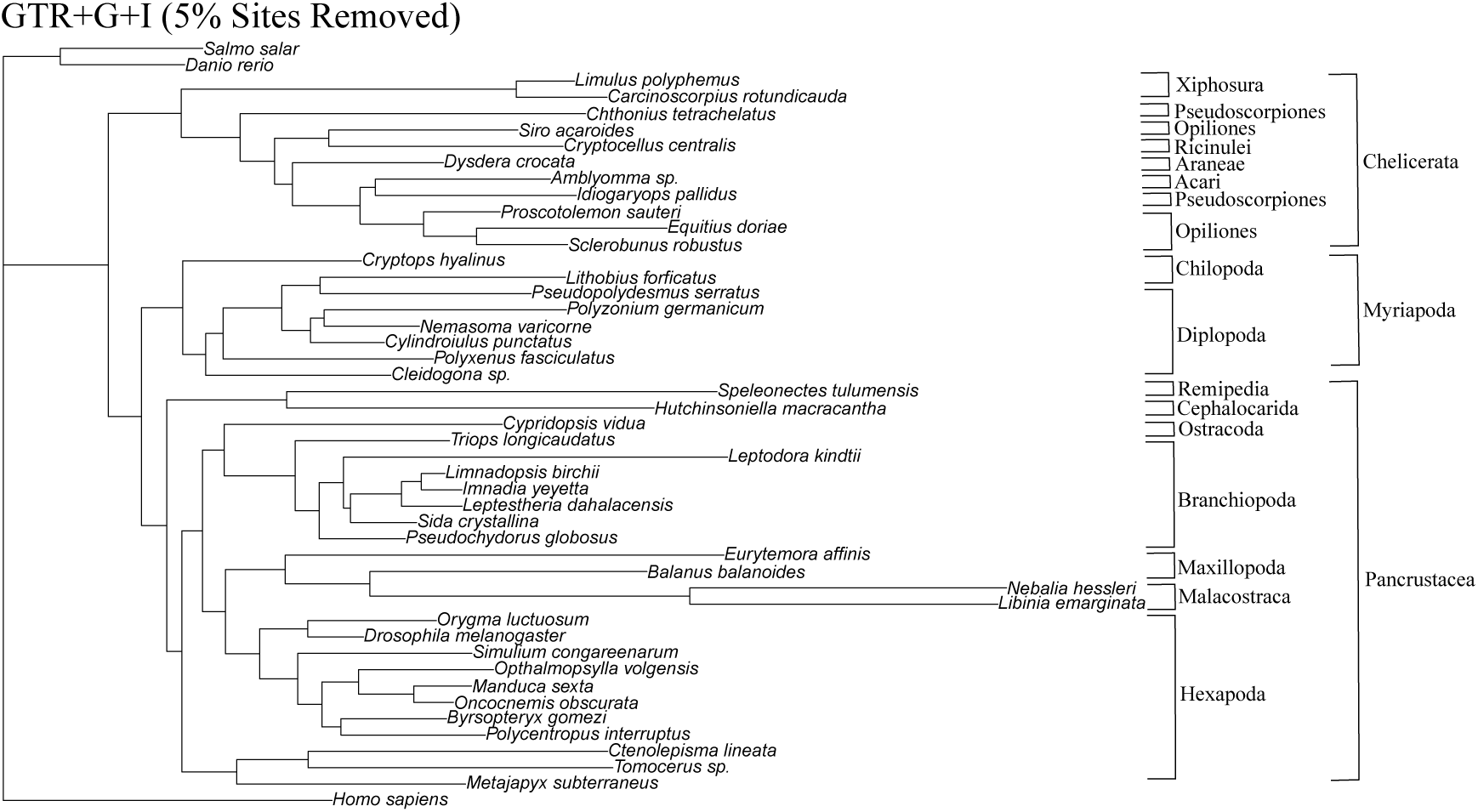
The tree inferred using the GTR+G+I model, with 5% of the sites removed from the arthropod dataset, represents the optimal tree among all those generated with this model.

**Figure S9:**
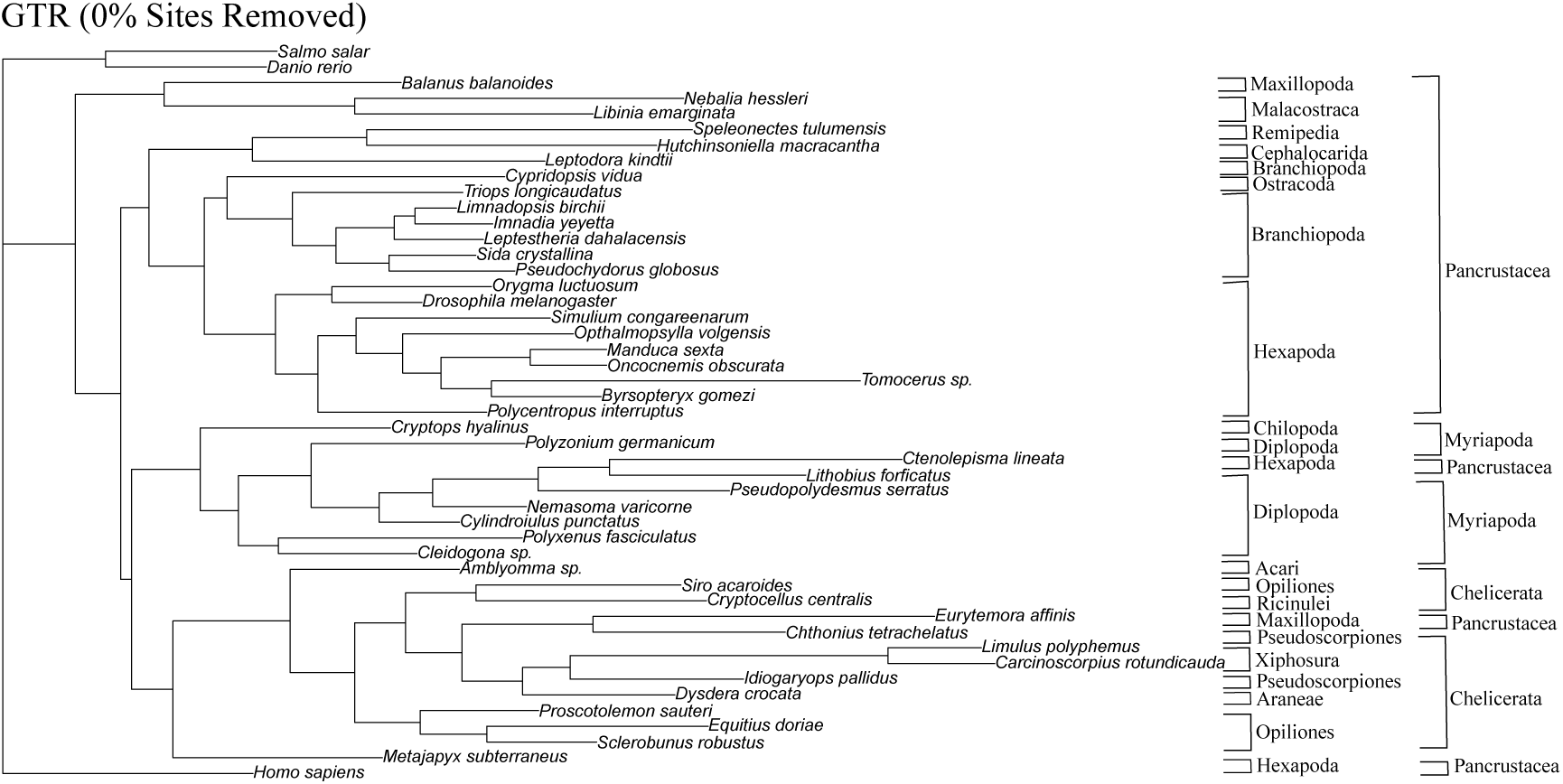
The tree inferred using the GTR model on the complete dataset.

**Figure S10:**
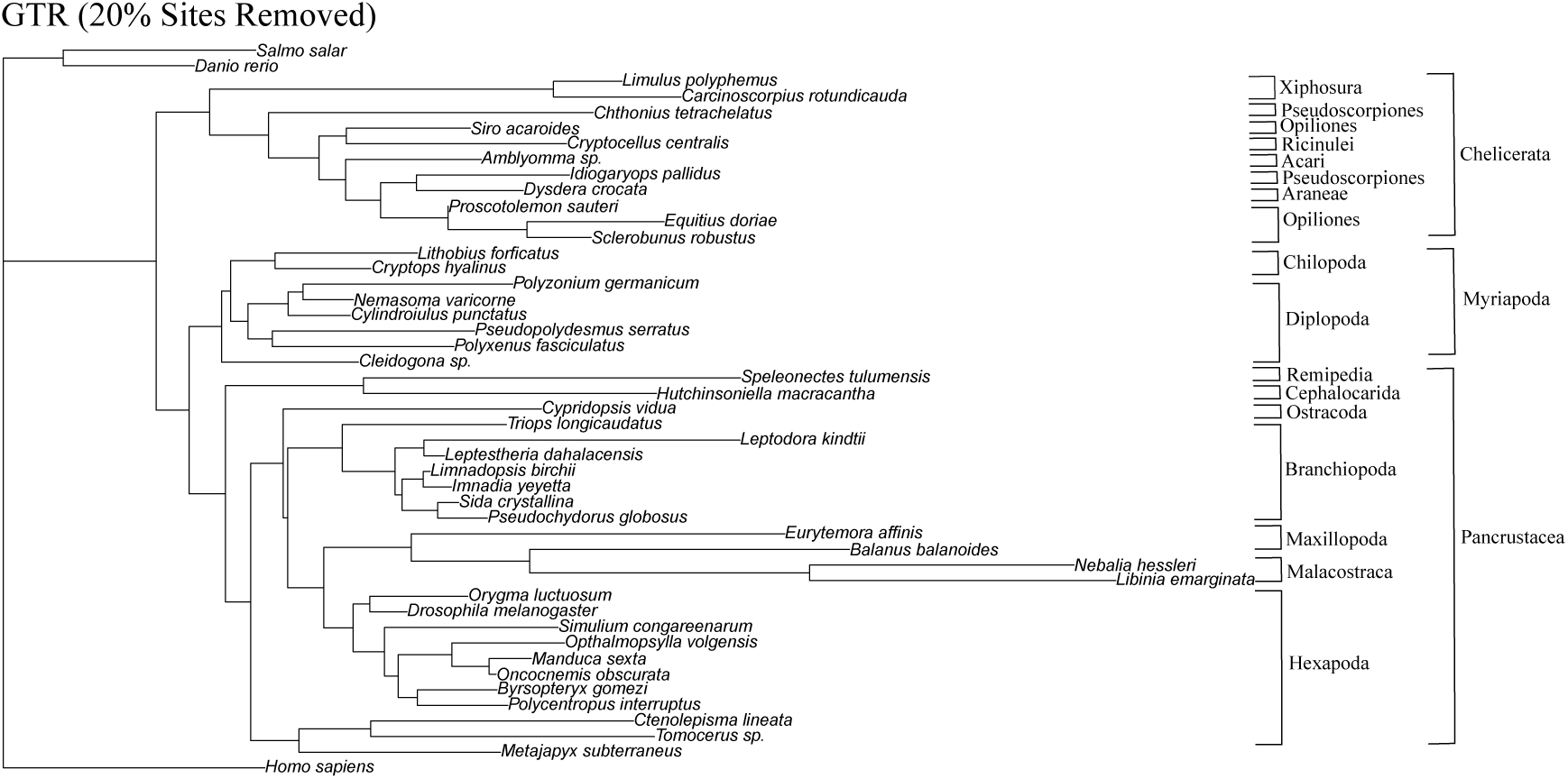
The tree inferred using the GTR model after removing 20% of the sites from the arthropod dataset. This is the optimal tree among those inferred with the GTR model.

